# Late- but not early-onset blindness impairs the development of audio-haptic multisensory integration

**DOI:** 10.1101/795500

**Authors:** Meike Scheller, Michael J. Proulx, Michelle de Haan, Annegret Dahlmann-Noor, Karin Petrini

## Abstract

Integrating different senses to reduce sensory uncertainty and increase perceptual precision can have an important compensatory function for individuals with visual impairment and blindness. However, how visual impairment and blindness impact the development of optimal multisensory integration in the remaining senses is currently unknown. Here we first examined how audio-haptic integration develops and changes across the life span in 92 sighted (blindfolded) individuals between 7 to 70 years of age by using a child-friendly size discrimination task. We assessed whether audio-haptic performance resulted in a reduction of perceptual uncertainty compared to auditory-only and haptic-only performance as predicted by maximum-likelihood estimation model. We then tested how this ability develops in 28 children and adults with different levels of visual experience, focussing on low vision individuals, and blind individuals that lost their sight at different ages during development. Our results show that in sighted individuals, adult-like audio-haptic integration develops around 13-15 years of age, and remains stable until late adulthood. While early blind individuals, even at the youngest ages, integrate audio-haptic information in an optimal fashion, late blind individuals do not. Optimal integration in low vision individuals follows a similar developmental trajectory as that of sighted individuals. These findings demonstrate that visual experience is not necessary for optimal audio-haptic integration to emerge, but that consistency of sensory information across development is key for the functional outcome of optimal multisensory integration.

**Research Highlights:**

- Audio-haptic integration follows principles of statistical optimality in sighted adults, remaining stable until at least 70 years of life
- Near-optimal audio-haptic integration develops at 13-15 years in sighted adolescents
- Blindness within the first 8 years of life facilitates the development of optimal audio-haptic integration while blindness after 8 years impairs such development
- Sensory consistency in early childhood is crucial for the development of optimal multisensory integration in the remaining senses

## Introduction

Early sensory input is crucial for the development of perceptual processes. A key method to discover the importance of early sensory input for perceptual development is to compare those who have had a sense, such as vision, impaired at an early developmental stage to those who acquire sensory deprivation later in life. For example, comparing humans who became blind early in life to those who became blind at older ages has revealed the impact of visual experience during development on other aspects of perception and cognition (Bedny et al., 2012; Pasqualotto, Furlan, Proulx, & Sereno, 2018; Wan et al., 2010a, see Scheller, Petrini, & Proulx, 2018 for a review). Reports on early blind individuals with extraordinary auditory or tactile abilities have nurtured the idea that non-visual perceptual mechanisms improve in order to compensate for the lack of visual information (Goldreich & Kanics, 2003; Gougoux et al., 2004a; Kolarik, Cirstea, & Pardhan, 2013; Norman & Bartholomew, 2011; Röder et al., 1999; Vercillo, Milne, Gori, & Goodale, 2015; Voss et al., 2004). For example, it has been shown that the brain of the early blind allows for changes in perceptual function through cortical reorganisation (Amedi, Raz, Pianka, Malach, & Zohary, 2003; Collignon et al., 2015; Ortiz-Terán et al., 2016). Several neuroimaging studies to date revealed structural and functional changes in the blind brain, such as increased fine-tuning of the auditory cortex (Huber et al., 2019), the redeployment of the visual cortex for non-visual tasks such as auditory localization and Braille reading (Gougoux, Zatorre, Lassonde, Voss, & Lepore, 2005; Sadato et al., 1996), or enhanced functional connections between uni-sensory and multisensory processing areas (Ortiz-Terán et al., 2016). These changes, together with enhanced auditory and tactile sensory functioning (Amedi et al., 2003; Collignon et al., 2013), support the hypothesis of cross-modal compensation. That is, the brain adaptively compensates for lacking visual input early during development, leading to enhanced non-visual perceptual functioning.

What several of these studies highlight is that the developmental time point of sensory deprivation determines how well an individual adapts to this perceptual state. That is, while congenitally blind individuals show enhanced auditory pitch discrimination or horizontal localisation abilities, late blind individuals do not exhibit such perceptual benefits (Gougoux et al., 2004b; Voss, Gougoux, Lassonde, Zatorre, & Lepore, 2006; Wan et al., 2010). Furthermore, studies on individuals that were born with dense bilateral cataracts, and who received sight-restoring treatment within the first months of life, showed that even a brief, transient phase of visual deprivation early in life leads to long lasting changes in visual and non-visual information processing (Collignon et al., 2015; Geldart, Mondloch, Maurer, De Schonen, & Brent, 2002; Guerreiro, Putzar, & Röder, 2016; Putzar, Hötting, & Röder, 2010; see Maurer, 2017 for a review). This stresses that sensory experience plays a critical role particularly during early developmental periods, when heightened cross-modal plasticity allows the individual to learn about the physical principles of the environment and their relation to their own body through sensory-motor contingencies (de Klerk, Johnson, Heyes, & Southgate, 2015; Nagai, 2019).

The sighted adult brain can integrate multisensory information by weighting the different sensory inputs by their reliability, in order to reduce sensory noise and increase perceptual precision and accuracy (e.g. Ernst & Banks, 2002; Rohde, van Dam, & Ernst, 2016). For example, while one can often easily hold a conversation without directly looking at a conversation partner (e.g. over the phone), this task becomes much more difficult when standing at a busy street. Here, visual information of the partner’s mouth movement can greatly enhance understanding of the conversation. However, the ability to optimally integrate sensory information has been found to only emerge late in childhood. While young children already possess the ability to make use of multisensory information (Neil, Chee-Ruiter, Scheier, Lewkowicz, & Shimojo, 2006), they do not perceptually benefit in the same way that adults do until 8-10 years of age (Adams, 2016; Gori, Sandini, & Burr, 2012), or even later (Nardini, Jones, Bedford, & Braddick, 2008; Petrini, Remark, Smith, & Nardini, 2014). For non-visual senses such as touch and sound, the developmental onset of optimal integration has not yet been established, but likely occurs after the age of 11 years (Petrini et al., 2014).

One prominent hypothesis, cross-modal calibration, accounts for this late development of optimal integration by suggesting that in early childhood the senses are kept separate to calibrate each other, thus impeding integration. During this time, the more robust sense for a certain task has been suggested to calibrate the less robust sense (Burr & Gori, 2012). For example, while touch is the more robust sense for estimating object size (Gori, Del Viva, Sandini, & Burr, 2008; Gori, Sandini, Martinoli, & Burr, 2010; Petrini et al., 2014), vision can be considered the more robust sense for estimating object orientation (Gori et al., 2008). In support of this hypothesis Gori and colleagues (2012) showed that haptic orientation discrimination performance is impaired in blind children because vision could not calibrate touch on this task (Gori, Tinelli, Sandini, Cioni, & Burr, 2012). Indeed, several other studies demonstrated that perceptual functioning in the remaining senses of blind individuals is severely compromised (Cappagli, Cocchi, & Gori, 2017; Cappagli, Finocchietti, Baud-Bovy, Cocchi, & Gori, 2017; Vercillo, Burr, & Gori, 2016; Zwiers, Van Opstal, & Cruysberg, 2001) when accurate performance depends on high resolution visual input (Coluccia, Mammarella, & Cornoldi, 2009; Gori, Sandini, Martinoli, & Burr, 2014; Pasqualotto et al., 2018; Pasqualotto & Proulx, 2012; Vercillo et al., 2016).

Most of the aforementioned studies on cross-modal compensation and cross-modal calibration assessed how visual impairment influences perception in the remaining, single senses. For example, Cappagli, Cocchi and Gori (2017) showed that early blind children and adults are severely compromised in the reproduction of hand pointing movements using proprioception, and struggle with extracting distance information from sound (Cappagli, Cocchi, et al., 2017). These findings show that unisensory processing in the remaining senses seems to depend on visual calibration early in development. However, much less is known about whether multisensory processes are affected by visual impairment in a similar way, although few studies tried to address this research question (Hötting and Röder, 2004; Champoux et al., 2011). However, it is still unknown how visual impairment affects optimal multisensory integration of the intact senses (e.g. audio-haptic optimal integration), and whether the onset and severity of visual impairment have a modulatory effect on it. As the visually impaired rely heavily on their remaining senses such as touch and hearing, it is crucial to understand when the ability to increase perceptual precision through optimal multisensory integration of the remaining senses is achieved. This knowledge would allow for the development of more effective sensory rehabilitation techniques that are functionally beneficial and meet the needs of the visually impaired individual (Ben Porquis et al., 2017; Gori, Cappagli, Tonelli, Baud-Bovy, & Finocchietti, 2016; Luo & da Cruz, 2016; Meijer, 1992, see Scheller, Petrini, & Proulx, 2018 for a review).

Here we used an optimised version of the audio-haptic size discrimination task from Petrini and colleagues (2014) to examine to what extent sighted and visually impaired adults and children reduce perceptual uncertainty by integrating sensory information from touch and hearing. We chose an object size discrimination task as haptic information tends to be the most robust sense for it, even in sighted children (Gori et al., 2008; Petrini et al., 2014) and thus should allow for an unbiased comparison that is not driven by differences in task difficulty and familiarity between the different vision groups. Based on the cross-modal compensation hypothesis, whereby intact senses compensate for impaired ones, an increased use of the non-visual senses would predict an earlier developmental onset of audio-haptic integration in low vision and blind individuals compared to sighted individuals. Furthermore, due to increased developmental plasticity early in life (Cappagli, Cocchi, et al., 2017; Collignon et al., 2013) we would predict that congenitally and early blind adults benefit more from integrating audio-haptic information, compared to late blind individuals. Based on the cross-modal calibration hypothesis, we would predict similar development of optimal audio-haptic integration in sighted, low vision, and blind individuals (independent of when vision was lost) as vision is not the most robust sense for this task and thus does not need to calibrate the other senses to achieve a more precise performance. Lastly, since recent findings (Cappagli, Finocchietti, Cocchi, et al., 2017; Cappagli, Finocchietti, Baud-Bovy, et al., 2017) have shown that children with low vision perform more similar to sighted than to blind children on different perceptual tasks, we predict that children and adults with low vision integrate audio-haptic information similar to sighted children and adults.

## Methods

### Participants

A total of 120 participants were recruited for this study. Of these, 46 were sighted adults (28 female, 41.6±18.2 years of age) and 46 sighted children (32 female, 11.5±2.5 years of age). They were grouped into five age groups in order to assess changes in multisensory integration over development. These age groups comprised of younger children (7-9 years), older children (10-12 years), adolescents (13-17 years), younger adults (18-44years), and older adults (45-70 years). For more details see Supplementary material S1.

Furthermore, three adults (two female, 30±16.8 years of age) and 11 children (six female, 10±2.1 years of age) with low vision, as well as nine totally blind adults (three female, 36±19 years of age), and five totally blind children (all male, 12.6±2.9 years of age) participated in the experiment. This sample size is similar to other studies assessing perceptual functioning in children and blind individuals (Cappagli et al., 2017; Garcia et al., 2015; Gori et al., 2010, 2014). Details of visually impaired (VI) participants are depicted in Table 1 and 2. The difference of interest between these groups is the presence or absence of visual experience during and after the first eight years of life, as this has been suggested to be the age at which vision-driven cross-modal calibration ends and children start integrating multisensory information in an adult-like fashion (Burr & Gori, 2012; Cappagli et al., 2017).

**Table 1:** Clinical and demographic information for blind and low vision adult participants.

| Participant | Sex | Age | Handedness | Age of Onset | Vision status | Diagnosis | Visual Acuity (Right Eye; Left Eye) [logMAR] | Vision group |
| --- | --- | --- | --- | --- | --- | --- | --- | --- |
| Vla1 | Female | 18 | Right | Birth | Congenitally Blind | Bilateral retinoblastoma, cataract, right enucleation | R - ; L = 2.8 | CB |
| Vla2 | Male | 59 | Right | Birth | Congenitally Blind | Glaucoma | R > 3; L > 3 | CB |
| Vla3 | Male | 21 | Right | Birth | Congenitally Blind | Congenital bilateral cataracts (until 9 years), Glaucoma, Retinal detachment |  | CB |
| Vla4 | Male | 33 | Right | 5.5 years | Early Blind | Glaucoma | R > 3; L > 3 | EB |
| Vla5 | Female | 18 | Right | 6 years | Early Blind | Retinitis pigmentosa | R > 1.8; L > 1.8 | EB |
| Vla6 | Female | 19 | Right | 7 years | Early Blind | Stargardt disease | R = 2.8; L = 2.8 | EB |
| Vla7 | Male | 60 | Right | 10 years | Blind | Leber's optic neuropathy | R = 1.5; L = 1.5 | LB |
| Vla8 | Male | 61 | Right | 11 years | Blind | Stargardt disease | R = 2.8; L = 2.8 | LB |
| Vla9 | Male | 35 | Right | 25 years | Blind | Macular degeneration, Retinopathy | R > 3; L = 2.8 | LB |
| Vla10 | Female | 49 | Right | 41 years | Low Vision | Pathological myopia, Choroidal neovascularization | R = 1.1; L = 0.8 | LV |
| Vla11 | Female | 19 | Right | Birth | Low Vision | Cataracts, Aniridia, Macular hypoplasia, Underdeveloped cornea | R = 1.1; L = 1.1 | LV |
| Vla12 | Male | 21 | Right | Birth | Low Vision | Ocular albinism, Nystagmus | R = 0.7; L = 0.7 | LV |

**Table 2:** Clinical and demographic information for blind and low vision child participants.

| Participant | Sex | Age | Handedness | Age of Onset | Vision status | Diagnosis | Visual Acuity (Right Eye; Left Eye) [logMAR] | Vision group |
| --- | --- | --- | --- | --- | --- | --- | --- | --- |
| Vlc1 | Male | 13 | Right | Birth | Blind | Bilateral microphthalmia, sclerocornea | R = 2.3; L = 2.3 | Blind (EB/CB) |
| Vlc2 | Male | 12 | Right | Birth | Blind | Retinal dystrophy; Leber's congenital amaurosis | R > 3; L > 3 | Blind (EB/CB) |
| Vlc3 | Male | 17 | ambi./right | 4 years | Blind | Retinal dystrophy | R > 3; L > 3 | Blind (EB/CB) |
| Vlc4 | Male | 9 | Left | 6 years | Blind | Glaucoma | R > 1.8; L > 3 | Blind (EB/CB) |
| Vlc5 | Female | 7 | ambi./right | Birth | Low Vision | Oculocutaneous albinism; Hypermetropia | R = 0.88; L = 0.76 | LV |
| Vlc6 | Male | 11 | Right | Birth | Low Vision | Red cone dystrophy | R = 0.7; L = 0.8 | LV |
| Vlc7 | Male | 12 | Right | Birth | Low Vision | Bilateral juvenile retinoschisis | R = 0.76; L = 1.3 | LV |
| Vlc8 | Female | 13 | Right | Birth | Low Vision | Stargardt disease | R= 1.0; L =1.0 | LV |
| Vlc9 | Male | 12 | Right | Birth | Low Vision | Cone dystrophy | R = 0.58; L = 0.94 | LV |
| Vlc10 | Male | 9 | Right | Birth | Low Vision | Stargardt disease | R = 1.0 ; L = 1.0 | LV |
| Vlc11 | Female | 7 | Right | Birth | Low Vision | Stargardt disease | R = 1.0; L = 1.0 | LV |
| Vlc12 | Female | 11 | Right | Birth | Low Vision | Stargardt disease | R = 1.04; L = 1.04 | LV |
| Vlc13 | Male | 11 | Right | 11 years | Low Vision | Neuromyelitis Optica | R = 1.5 L = 0.3 | LV |
| Vlc14 | Female | 9 | Right | 3.5 years | Low Vision | Bilateral optic atrophy and nystagmus | R = 1; L = 1.3 | LV |
| Vlc15 | Female | 8 | Right | 4 years | Low Vision | Stargardt disease | R = 0.4; L = 0.3 | LV |
| Vlc16<sup>†</sup> | Male | 12 | Right | Birth | Low Vision | Congenital Glaucoma, Left enucleation | R = 1.1; L = - | Blind (EB/CB) |

<sup>†</sup>data from this individual could not be used

All participants had normal hearing and no other certified developmental disorders, such as Autism Spectrum Disorder. Data from one blind child (VIc16) was excluded from the analysis due to inability to pay attention and complete the task due to hyperactive behaviour, leaving data of four blind children. Handedness was assessed using the Oldfield Edinburgh Handedness Inventory (Oldfield, 1971). All adults and parents of sighted and visually impaired children gave informed consent before participating in the study, which received ethical approval from the University of Bath Ethics Committee (ref # 15-211) and the National Health Research Authority (IRAS ref # 197917). Sighted adults and children were recruited through local schools, University advertisements, and Research Participation Panels. Visually impaired individuals were recruited through Moorfields Eye Hospital, Bristol Eye Hospital, local charities for the visually impaired, word of mouth, and University advertisements.

### Stimuli

Stimuli development was based on a standardised and validated method by Petrini et al. (2014). The stimuli consisted of 17 white, 3D-printed plastic balls of different sizes, ranging from 41mm to 57mm in diameter with an increment size of one millimetre. The median ball size with a diameter of 49mm was chosen as standard stimulus, leaving eight comparison stimuli bigger than the standard ball (50mm-57mm) and eight smaller comparison stimuli (41mm-48mm). A sound recorded from the standard ball with 49mm diameter was used to create the comparison balls sound. Praat software (Boersma, 2001) was used to modulate the sound in amplitude to match the sizes of all comparison balls, resulting in sixteen comparison sounds ranging from 71dB to 79dB. The increment size for auditory stimuli was 0.5dB and has been matched to the haptic stimuli in accordance with Petrini et al. (2014), in which 2mm haptic size increment were used with 1dB sound amplitude increments. Pilot tests confirmed the audio-haptic stimulus pair to be well adjusted.

### Procedure

The participant was blindfolded and seated comfortably in a chair in front of a table and was blindfolded in order to eliminate any visual cues during the experiment. The set up on the table comprised of a touch screen panel on which the haptic stimuli (plastic balls) were placed during the experiment, one at a time (see Fig. 1). A thin layer of foam between the ball and touch screen prevented the stimuli from generating impact sounds when being placed down. The participant’s dominant hand rested on a soft foam block, which was positioned next to the touch screen. During each trial, a ball was placed on the touch screen in front of the participant, who was then asked to briefly tap the ball with the straight and flat palm of their dominant hand. As the participant was blindfolded, their hand was guided by the experimenter. Once pressure was sensed on the touch screen the corresponding sound, which provided the auditory size information, was played back through headphones. After tapping the ball, the hand was returned to the soft foam block and the same procedure was repeated with a second stimulus. After two stimuli (unimodal) or two stimuli-pairs (bimodal) were presented, the participant was asked to indicate whether the first or the second object was bigger. Before each experimental block (condition), participants received training on at least four practice trials in order to indicate whether they were able to do the task and to familiarize them with the stimuli.

**Figure 1.**
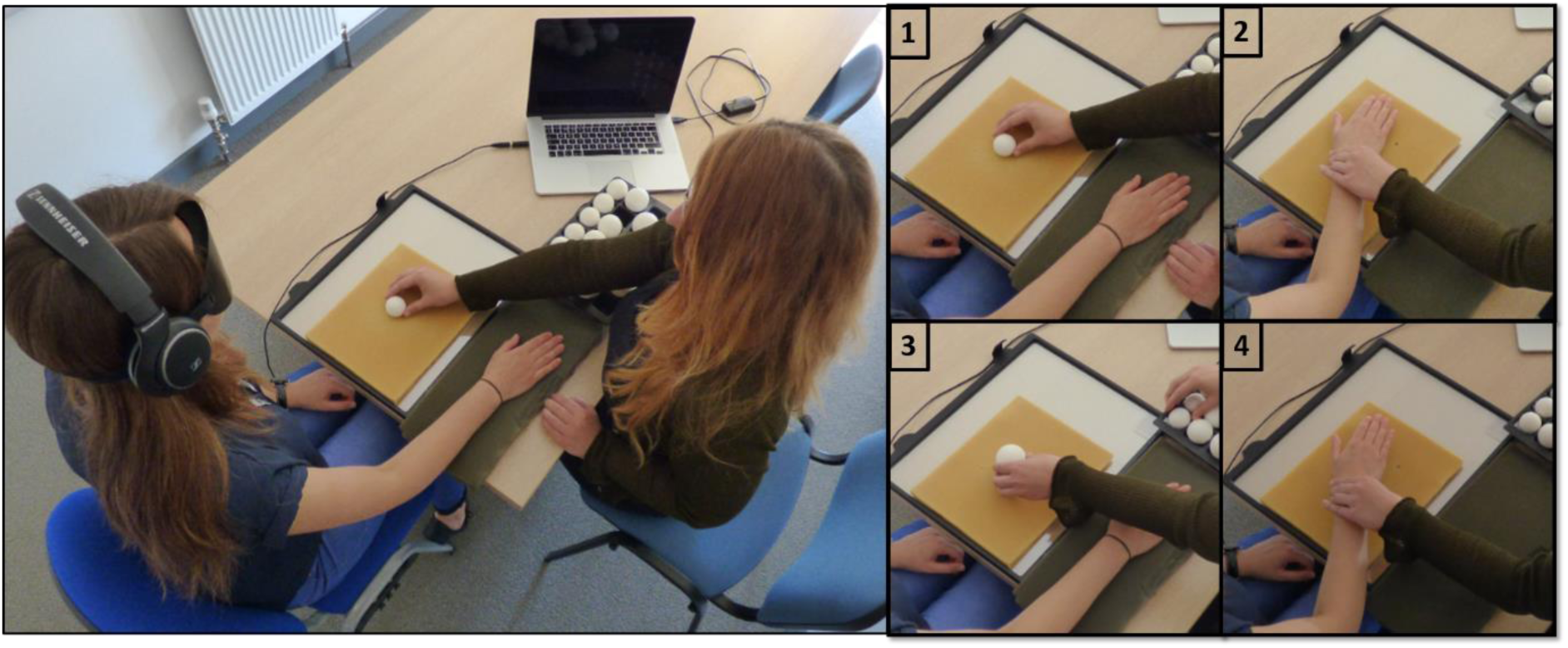
Experimental set up and procedure. All participants were blindfolded and sat in front of the set up with their dominant hand resting on a semi-soft foam surface. (1) Haptic stimuli were positioned in a pre-defined location on a thin foam surface that was placed on a touchscreen in front of the participant. (2) Their dominant hand was guided to the location of the stimulus, which they briefly tapped with the flat and straight hand. In the haptic condition, only information from touch was available. In the bimodal conditions, the pressure that was sensed by the touch screen elicited the size-corresponding sound to be played back through headphones. In the audio only condition, participants held a pen, which they used to tap on the touch screen to trigger the sound. In this condition their hand was guided as well. (3,4) The same procedure repeated for a second stimulus. Participants were then asked to judge which of the two objects was bigger.

During each trial, the standard stimulus (49mm ball, 75dB sound) was compared to either a bigger or a smaller stimulus. The order in which standard or comparison stimuli were presented was random – with the standard being either first or second. The following stimulus conditions were grouped into blocks of 30 trials in a counter-balanced order: (a) audio only, (b) haptic only, (c) bimodal congruent, and (d) bimodal incongruent. In the audio-only condition, participants only discriminated between object sizes based on the sounds they heard through headphones. Sounds were triggered by participants tapping on the touch screen with a pen. Their hand was guided by the experimenter in order to match the timing of arm movement in the other blocks. Triggering the sound through tapping was used to allow comparison between blocks that all used active arm movement and to control for attentional shift due to expected sound onset. In the haptic only condition, participants tapped the ball, but the sound was not played back. Bimodal congruent presentations played the corresponding sound when the ball was tapped. In the bimodal incongruent condition sound and touch gave conflicting size information. i.e. a bigger ball (53mm) was presented with the sound of a smaller ball (73dB = 45mm), together averaging on the standard stimulus size (49mm). This cross-modal conflict between haptic and auditory information can be used to determine the degree of perceptual bias towards one of the two cues, and with that the relative reliability (or attributed weight) of the two modalities for this task. We used only one incongruent condition, as Petrini et al. (2014) reported no differences between incongruent pairings. Limiting the length of the experiment is especially important with respect to testing children and individuals with shorter attention spans. Responses were used to calculate discrimination thresholds for each condition, which serve as a measure for perceptual precision. Lower discrimination thresholds indicate a higher perceptual precision. For further information on the procedure and data analysis, see the Supplementary material S2.

## Results

Size discrimination thresholds were used as a measure of precision and were estimated for all participants and conditions separately. All data were assessed for normality, homogeneity of variances and outliers before appropriate tests were chosen. Test assumption checks are reported in the Supplementary material S3.

To assess how size discrimination thresholds for audio, haptic, and audio-haptic stimuli differ between age groups we carried out a mixed factorial ANOVA, using the three conditions as within-subjects factor and age group as between-subjects factor. The analysis indicated significant main effects for age (*F*(4,87) = 8.975, *p* < .001) and condition (*F*(2,174) = 12.93, *p* < .001), as well as a significant interaction between age group and condition (*F*(8,174) = 2.856, *p* = .005). Bonferroni-corrected, paired t-test were used to compare discrimination thresholds between age groups. Below, we report corrected p-values. Effect sizes were computed as Hedges *g* with correction for small sample sizes (*d_unbiased_*, Cumming, 2012). Younger adults performed significantly better in the audio-haptic bimodal condition than with either auditory (*t*(29)= 4.85, *p* < .001, *d_unb_* = 0.874) or haptic (*t*(29) = 2.28, *p* = 0.015, *d_unb_* = 0.411) information alone. Similarly, the older adults performed significantly better in the bimodal condition than in either the auditory (*t*(14)= 4.06, *p* = .002, *d_unb_* = 1.018) or haptic (*t*(14)= 4.10, *p* = .002, *d_unb_* = 0.703) condition. In both, the young and older children groups, thresholds in the bimodal condition did not differ from either the auditory-only (7-9yo: *t*(7) = 0.239, *p* = 1, *d_unb_* = 0.153; 10-12yo: *t*(21) = 1.15, *p* = .394, *d_unb_* = 0.241) nor haptic-only (7-9yo: *t*(7) = 0.45, *p* = 1, *d_unb_* = 0.203; 10-12yo: *t*(21) = 2.32, *p* = 1, *d_unb_* = 0.485) condition. In adolescents, bimodal discrimination thresholds were significantly lower than in the auditory-only condition (*t*(14) = 3.01, *p* = .014, *d_unb_* = 0.756), but only marginally lower than in the haptic-only condition (*t*(14) = 2.32, *p* = .054, *d_unb_* = 0.584) condition. The results are depicted in Fig. 2, showing a clear trajectory of the improvement of size discrimination performance with age. In order to compare discrimination performance in the multisensory condition with Bayes-optimal integration performance, we calculated predictions for discrimination thresholds based on maximum likelihood estimation (MLE, see equation 1.3 and 1.4 in Supplementary material S2) for each individual separately. Averages for predicted bimodal thresholds are depicted in Fig. 2 as black circles. For more details on individual integration performance see Supplementary material S4. Comparing the bimodal threshold to MLE prediction, we found that only in the two adult groups discrimination thresholds did not differ from MLE prediction (18-44 year-olds: *t*(29) = 2.1, *p* =.133, *d_unb_* = 0.379; 45-70 year-olds: *t*(14)= 0.94, *p* = 1, *d_unb_* = 0.229). Sensory weights for auditory and haptic cues indicated that all groups, apart from older adults, weighted haptic cues stronger than auditory cues. For more details on cue weighting see Supplementary material S5.

**Figure 2.**
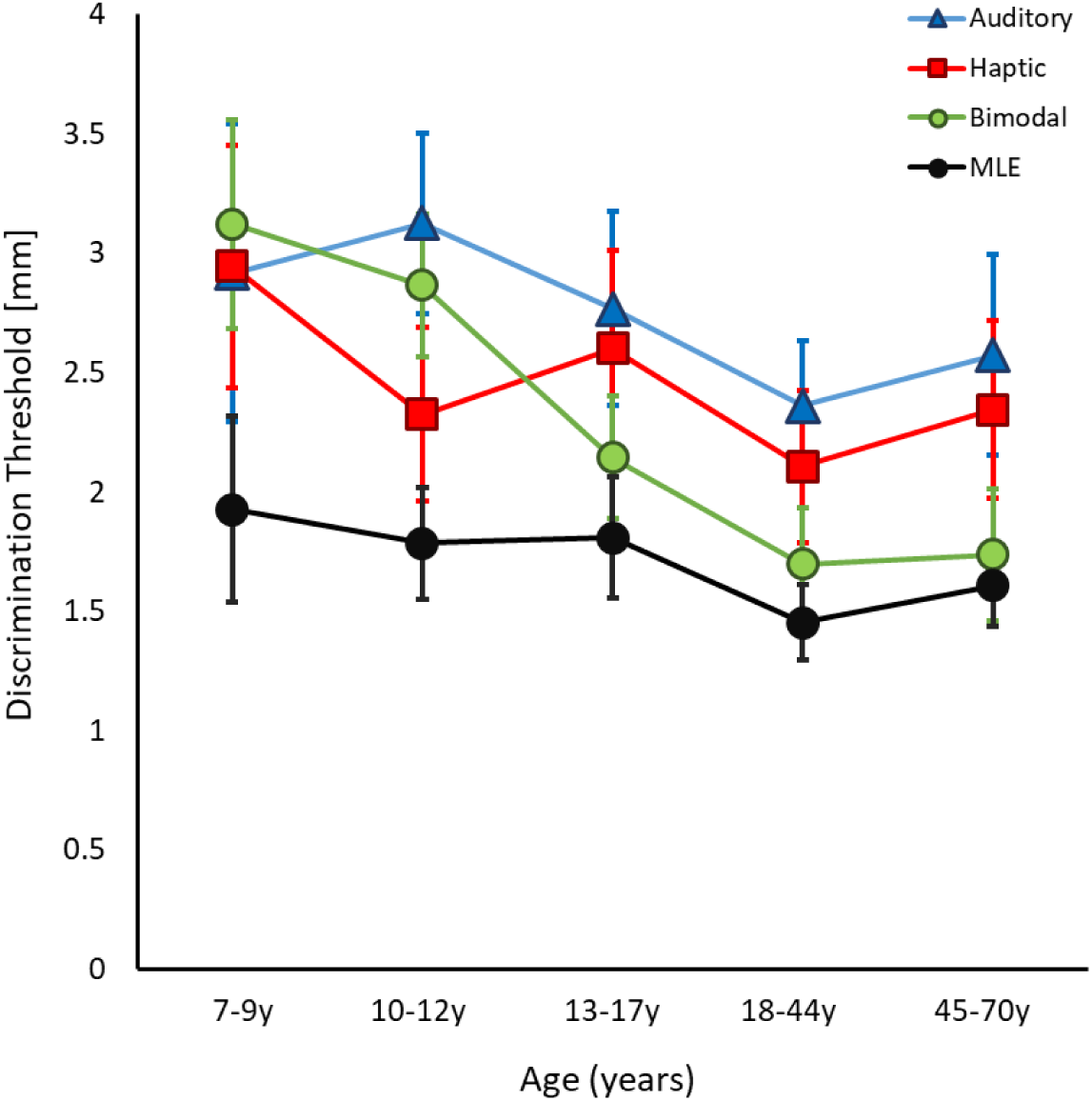
Unimodal and bimodal discrimination thresholds of sighted individuals of different ages. Average size discrimination thresholds for all conditions across five age groups. Measured discrimination thresholds for auditory-only (blue triangles), haptic-only (red squares), as well as bimodal (green circles) conditions plotted for five age groups, including younger children, older children, adolescents, as well as younger and older adults. Black circles represent the average discrimination thresholds predicted by Bayed optimal prediction (MLE) and were calculated as a weighted combination of the two unimodal estimates for each individual. Error bars represent 95% CIs.

To examine the extent to which the development of audio-haptic integration depends on visual input, we assessed audio-haptic discrimination performance in adults and children with different levels of visual experience. Thereby we focused on individuals with reduced visual input (low vision, logMAR < 1.3, *n* = 15) and no functional visual input (blind, logMAR ≥ 1.3; *n =* 14) separately. The grouping was based on the WHO definition of blindness using individual visual acuity measures (World Health Organization, 2018). In the low vision group, integration performance was compared between adults and children to assess whether a reduction in visual input affects how audio-haptic integration develops. To assess how the absence of vision and the developmental time point of vision loss affect audio-haptic integration, we then compared integration performance between blind adults with three different onsets of vision loss: congenitally blind, early blind, and late blind. We chose eight years as a developmental cut-off age to differentiate between early and late blind, as this has been identified as the earliest age at which adult-like multisensory integration emerges in sighted children when using vision (Adams, 2016; Gori et al., 2008; Nardini et al., 2008, see Fig. 9 in discussion). Furthermore, it has been proposed that vision-driven cross-modal calibration takes place within the first eight years of life (Cappagli et al., 2017). In cases where both eyes were affected differently (e.g. participant VIc13) the visual function of the better eye was used as an approximation of best visual function. Non-parametric tests were applied for all analyses including visually impaired individuals as the sample size was small in all sub-groups. Bonferroni-corrected Mann-Whitney U tests were used for group comparisons, while Crawford-Howell case-control comparisons (Crawford, Garthwaite, & Porter, 2010) were used for individual performance comparisons.

The influence of reduced visual input on audio-haptic integration was examined by comparing discrimination thresholds of children and adults with low vision against the respective developmental group of sighted participants. Comparing low vision children (aged 7-12 years) with sighted children (aged 7-12 years) showed that discrimination thresholds did not significantly differ between groups in neither auditory-only (*U* = 151, *p* = 1, *r* = 0.04), haptic-only (*U* = 158, *p* = 1, *r* = 0.10), nor audio-haptic (*U* = 171, *p* = 1, *r* < 0.01) conditions. Furthermore, there was no difference between adults with low vision and adults with typical sight in either condition (auditory: *U* = 74, *p* = 1, *r* = 0.04; haptic: *U* = 39, *p* = .674, *r* = 0.18; audio-haptic: *U* = 43, *p* = .890, *r* = 0.15, see Fig. 3).

**Figure 3:**
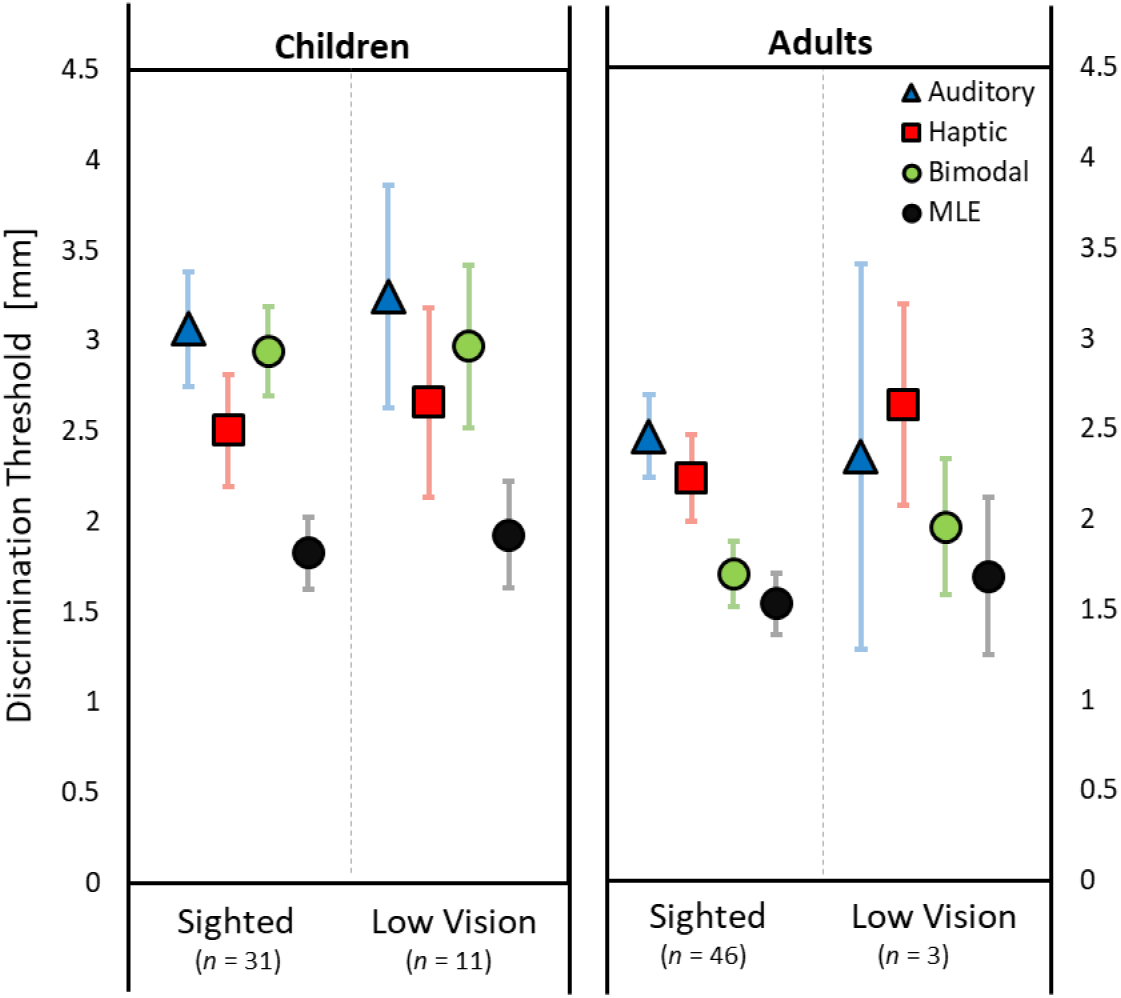
Unimodal and bimodal discrimination thresholds of sighted and low vision individuals. Average size discrimination thresholds of both unimodal and bimodal conditions, as well as Bayes optimal prediction (MLE). Left panel shows average thresholds for children, while the right panel shows discrimination thresholds for adults, with the sighted group averages plotted as reference. Error bars represent 95% CIs.

The influence of functional visual input on audio-haptic integration was assessed by comparing discrimination thresholds of typically sighted children and adults to that of blind children and adults with different onsets of blindness (congenitally, early, and late blind). Each individual blind child was compared to the respective age group described in the sighted section above (7-9years, 10-12years, 13-17 years) using Crawford-Howell t-tests for case-control comparisons. Most comparisons did not reach significance (*p* > .05), however, the 9-year old early blind child showed a significantly lower discrimination threshold only in the bimodal condition, compared to sighted 7-9 year olds (*t* = 3.47, *p* = .025, zC*C* = 3.66, see Fig. 4).

**Figure 4:**
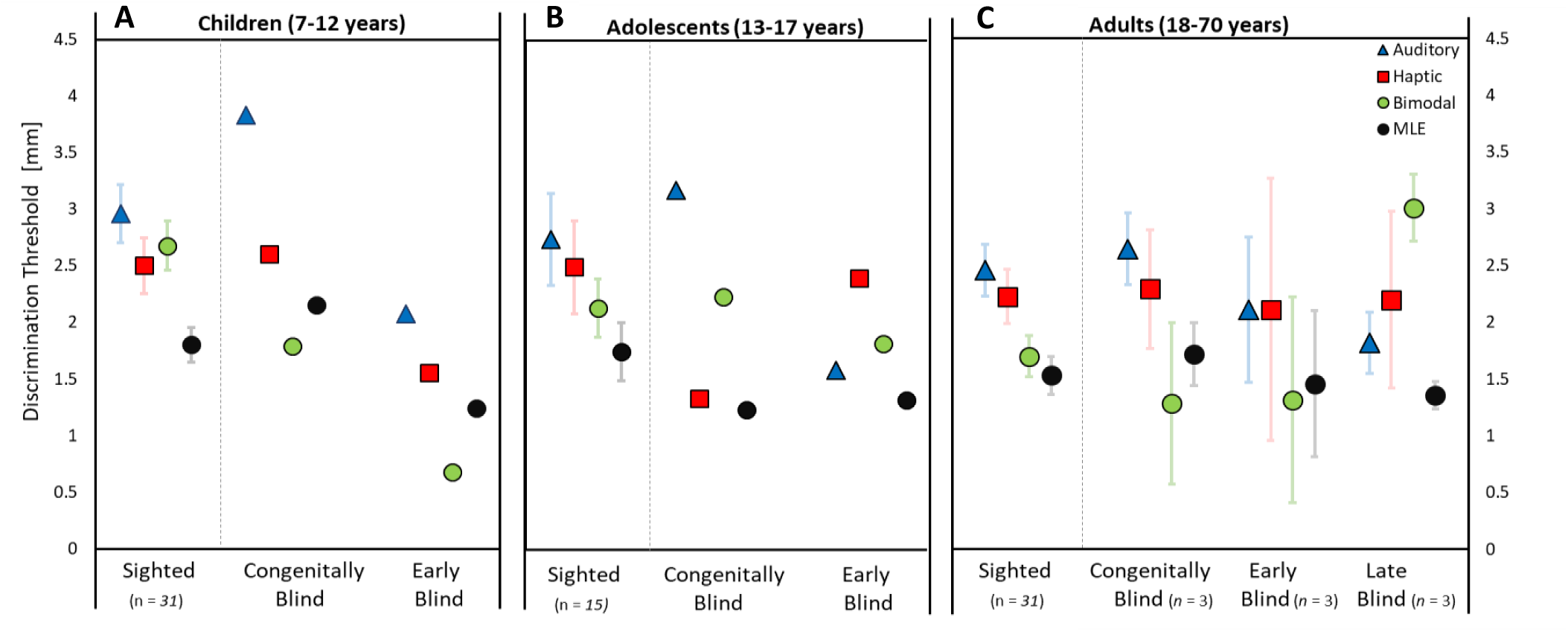
Discrimination thresholds for unimodal and bimodal performance for sighted and blind individuals. Average size discrimination thresholds for both unimodal and bimodal conditions, as well as Bayesian model prediction (MLE). Panel A shows thresholds for two blind children aged 9 and 12, as well as the average thresholds for children aged 7-12 years. Panel B shows thresholds for two blind adolescents aged 13 and 17, together with the average thresholds for 13-17-year-old sighted adolescents. Panel C shows average thresholds for adults with congenital, early, or late blindness onset, as well as the sighted adult thresholds for reference on the left. Early blindness is defined as having an onset within the first 8 years of life, while late blindness is defined by an onset after 8 years of life, in line with the duration of cross-modal calibration (Burr & Gori, 2012). Black circles represent the average discrimination thresholds predicted by maximum likelihood estimation based on a weighted combination of the two unimodal estimates. Error bars represent 95% CIs.

Discrimination thresholds of blind adults were assessed, similar to low vision adults, on the basis of group comparisons using Bonferroni-corrected Mann-Whitney U-tests. There were no significant differences between the congenitally blind, nor the early blind individuals and sighted adults in either the auditory (CB: *U* = 47, *p* = 1; EB: *U* = 83, *p* = 1), haptic (CB: *U* = 35, *p* = .499; EB: *U* = 67, *p* = 1), or audio-haptic conditions (CB: *U* = 91, *p* = .951; EB: *U* = 82, *p* = 1). However, the late blind individuals differed from sighted adults in the audio-haptic condition, showing higher discrimination thresholds (*U* = 9, *p* = .038, *r* = 0.36, see Fig. 4), while they did not differ in either auditory (*U* = 108, *p* = .254) or haptic thresholds (*U* = 91, *p* = .951).

### Multisensory benefit (Δ_measured-predicted_)

We next computed the differences between bimodal discrimination thresholds and MLE predictions Δ_measured-predicted_ for each individual. This measure provides a quantified estimation of the perceptual benefit that is gained through multisensory integration. Differences between bimodal threshold and MLE prediction across the developmental age range are depicted for sighted individuals in Fig. 5, and for low vision and blind individuals in Fig. 7.

**Figure 5.**
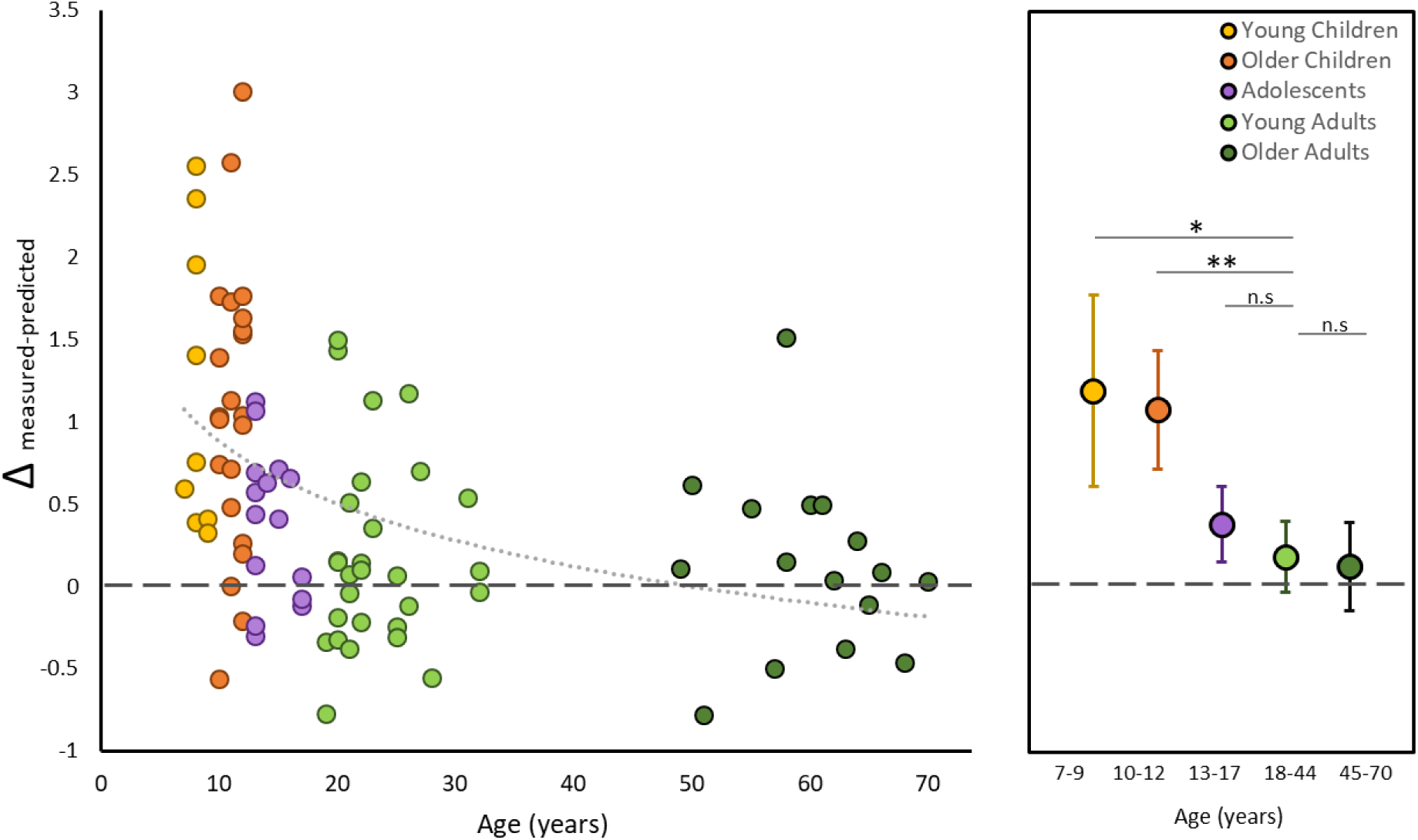
Integration performance of typically sighted individuals. Left panel shows individual threshold differences between predicted and measured discrimination thresholds for audio-haptic bimodal stimulus presentation across all ages. The dashed line at y = 0 indicates optimal performance predicted by MLE, which is based on the auditory and haptic unisensory estimates. Data below this line indicates an increase in precision that is better than predicted by the model. Different colors correspond to the different age groups: young children (7-9 years), older children (10-12 years), adolescents (13-17 years), younger adults (18-44 years), and older adults (45-70 years). Light grey trend line indicates the line of best fit. The right panel shows means for discrimination threshold difference scores (Δ) for each age group separately. Error bars indicate 95% CI. * = p<.05; ** = p< 0.01; n.s. = not significant.

Comparing the multisensory benefit Δ_measured-predicted_ of young adults with the different developmental age groups, we found young adults and older adults did not differ from each other (*t*(29) = 0.33, *p* = 1 *d_unb_* = 0.101). Furthermore, the multisensory benefit of adolescents aged 13-17 years did not differ from that of young adults either (*t*(35) = 1.23, *p* =.568, *d_unb_* = 0.357). Contrastingly, older children as well as young children significantly differed from young adults in the perceptual benefit gained through multisensory integration (7-9yo: *t*(9) = 2.81, *p* = .039, *d_unb_* = 1.319; 10-12 yo: *t*(35) = 4.19, *p <* .001, *d_unb_* = 1.231; see Fig. 5).

In order to assess how integration performance develops in low vision individuals we compared the multisensory benefit Δ_measured-predicted_ between sighted and low vision children, and between sighted and low vision adults. Average scores were not significantly different between sighted and low vision individuals, this was true for both children (*U* = 158, *p* = .735, *r* = 0.10) and adults (*U* = 83, *p* = .543, *r* = 0.02, see Fig. 6).

**Figure 6.**
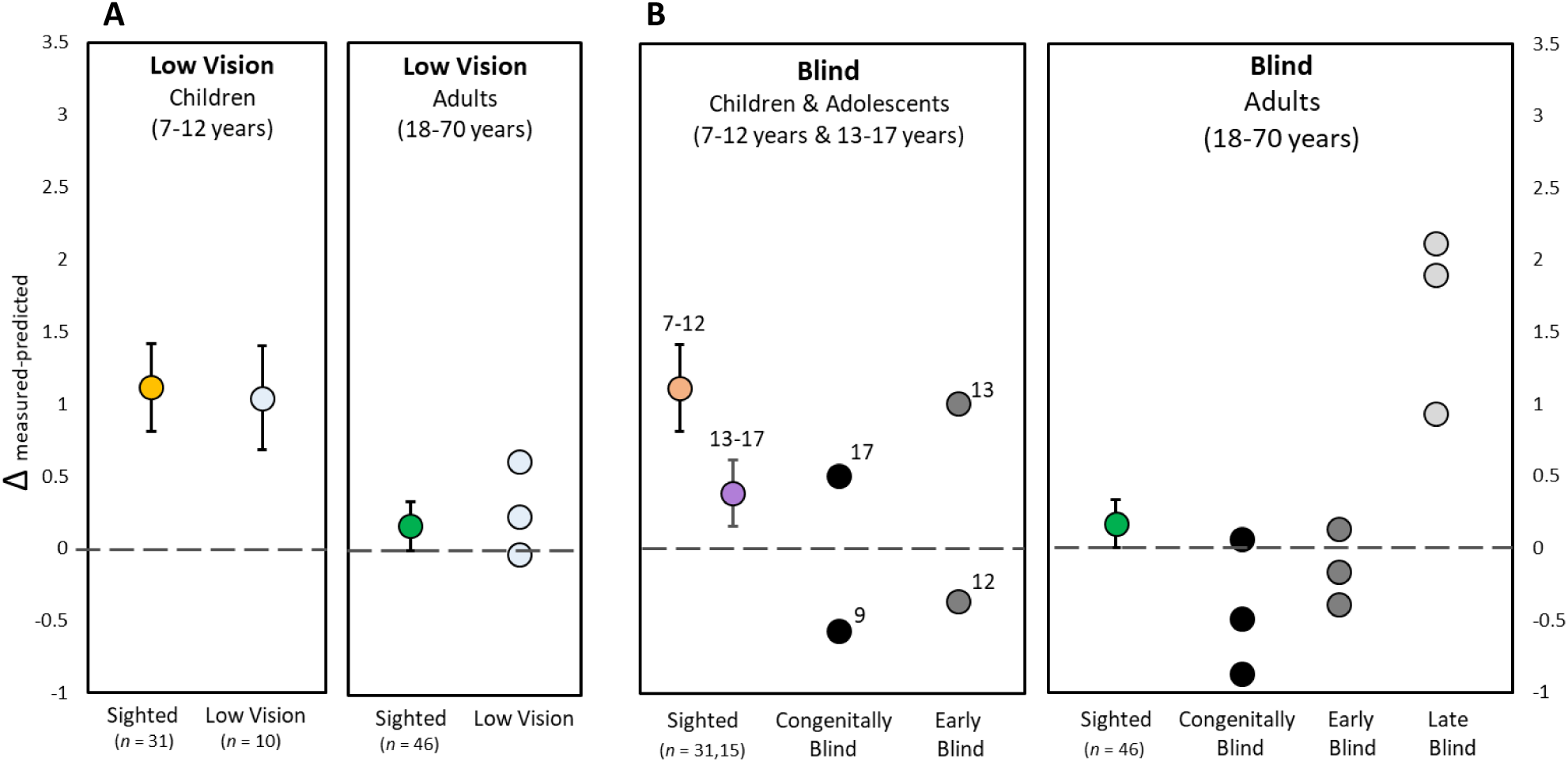
Integration performance of low vision and blind individuals. Differences between predicted and measured discrimination thresholds for bimodal stimulus presentation. A shows average multisensory benefit scores for children, and individual multisensory benefit scores for adults with low vision (light circles). The average multisensory benefit scores of the respective, age-matched sighted groups are plotted as references. B shows multisensory benefit for individual congenitally blind (black), early blind (grey), and late blind (light grey) individuals. Early and late blindness are defined by the onset of blindness either before or after the age of 8 years. For the children and adolescents, individual ages and age ranges are indicated next to the data points to allow for a direct comparison. The dashed line at y = 0 indicates MLE model prediction based on the auditory and haptic unisensory estimates, while data below this line indicates an increase in precision that is better than predicted by the model. Error bars indicate 95% CI.

Comparing the average Δ_measured-predicted_ between individual blind children and the age-matched sighted children (7-12years) or adolescent (13-17years) groups indicated that the congenitally blind 9-year old benefitted from integrating audio-haptic information significantly more than sighted children (*t* = 1.92, *p* = .032, zC*C* = 1.96). For the 12-year old early blind individual, there was a marginal difference (*t* = 1.69, *p* = .051, z*CC* = 1.72, suggesting that this individual also reduced uncertainty more than sighted children. We did not find any differences between the 17-year old congenitally blind individual and sighted adolescents (*t* = 0.25, *p* = .105, zC*C* = 1.36), nor for the 13-year-old early blind individual and sighted adolescents (*t* = 0.25, *p* = .403, zC*C* = 0.26). Next, we compared sighted adults with blind adults in three different blindness onset groups (congenitally, early, late blind). Congenitally blind individuals integrated audio-haptic information optimally, or even super-optimally (see Fig. 6). This group differed from sighted adults only marginally (*U* = 112, *p* = .059, *r* = 0.24). Discrimination thresholds of early blind individuals did not differ significantly from that of sighted adults (*U* = 92, *p* = .322, *r* = 0.07). Lastly, late blind individuals showed significantly higher Δ_measured-predicted_ scores compared to sighted individuals (*U* = 5, *p* = .002, *r* = 0.448), indicating reduced integration performance. Fig. 4 shows late blind adults exhibit similar auditory and haptic thresholds as other adults. Differences between bimodal threshold and MLE prediction for blind children and adults, as well as the respective sighted age groups, are depicted in Fig. 6. For an overview of individual scores for adults and children from all vision groups across the developmental age range see Fig. 7.

**Figure 7.**
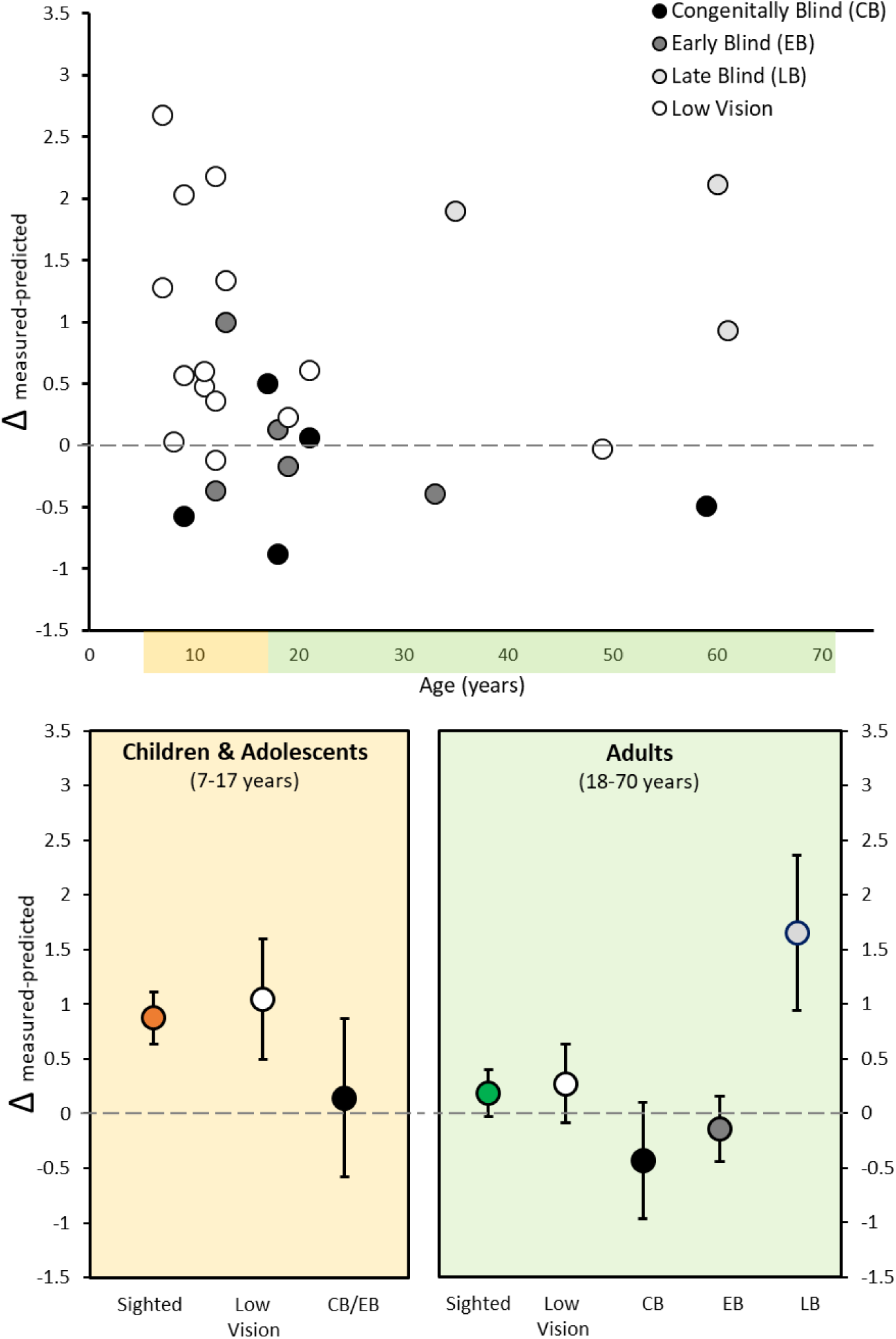
Overview of multisensory integration performance in low vision and blind individuals. Panel A shows individual difference scores for measured versus predicted discrimination thresholds as a function of age. Predicted threshold, indicated by the grey dashed line at y = 0, is based on the Bayesian integration model predicting optimal integration performance. Model predictions have been calculated for each participant separately and are based on the individual auditory and haptic unisensory thresholds. Individuals are color-coded based on different amounts of visual experience. Panel B shows average scores for low vision and blind children and adolescents, with the sighted children and adolescent group as a reference. Panel C shows average scores for low vision and blind adults, depending on the age of blindness onset. Average scores for sighted adults are plotted as a reference. Error bars indicate 95% CI.

## Discussion

The brain’s ability to enhance perceptual precision by integrating input from multiple senses develops late in sighted individuals (Adams, 2016; Gori et al., 2008; Nardini et al., 2008; Petrini et al., 2014). Early blindness has been shown to impact on non-visual perception in two ways: on the one hand, neural plasticity allows the individual to cross-modally compensate for missing sensory input, for example through enhanced tactile discrimination or auditory localisation (Amedi et al., 2003; Collignon et al., 2013). On the other hand, blindness precludes the calibration of the non-visual senses through vision. This has been shown to lead to impaired auditory or proprioceptive spatial perception (Cappagli et al., 2017; Gori et al., 2014). However, as most of our environment is multisensory, and as visually impaired individuals rely more heavily on other senses such as touch and hearing, the functional outcomes of visual deprivation on the benefits of audio-haptic integration (reducing sensory uncertainty by combining sensory information) are of fundamental importance.

Here we report, for the first time, that while congenitally and early blind (EB) adults show similar or even marginally better integration performance than sighted adults, audio-haptic integration performance of late blind adults is impaired. As expected, the developmental period during which visual experience influences the development of audio-haptic integration extends until eight to nine years of life. This falls in line with the previously proposed period of cross-modal calibration through vision (Cappagli et al., 2017; Gori et al., 2014). Based on the idea that during development the more robust sense calibrates the less robust senses, we would expect that the presence or absence of visual experience would not affect the performance on our audio-haptic size discrimination task. This is because touch is the more robust sense for assessing size information, compared to audition (Petrini et al., 2014; *present study*) or vision (Gori et al., 2008, 2012). Indeed, we find that blindness early in life does not affect audio-haptic integration later in life, which would therefore support the idea that the more robust sense teaches the less robust sense and that vision is not necessary for audio-haptic integration. However, we also find that early blindness seems to lead to an earlier development of optimal audio-haptic integration. This finding would support the idea of cross-modal compensation. That is, an increased use of the remaining senses leads to an enhanced recruitment of presumptive “visual” areas in the brain to process non-visual information, thereby enhancing performance in those senses (Amedi et al., 2003; Collignon et al., 2013). However, in contrast to both these theories we find that late blindness, which indicates the presence of visual experience during early development, leads to a disruption in audio-haptic integration performance. While the presence of visual experience early in life reduces audio-haptic integration performance in the late blind, it does not reduce integration performance in the sighted. These findings cannot be explained by either cross-modal calibration or sensory compensation alone. A summary of these findings can be seen in Fig. 8.

**Figure 8.**
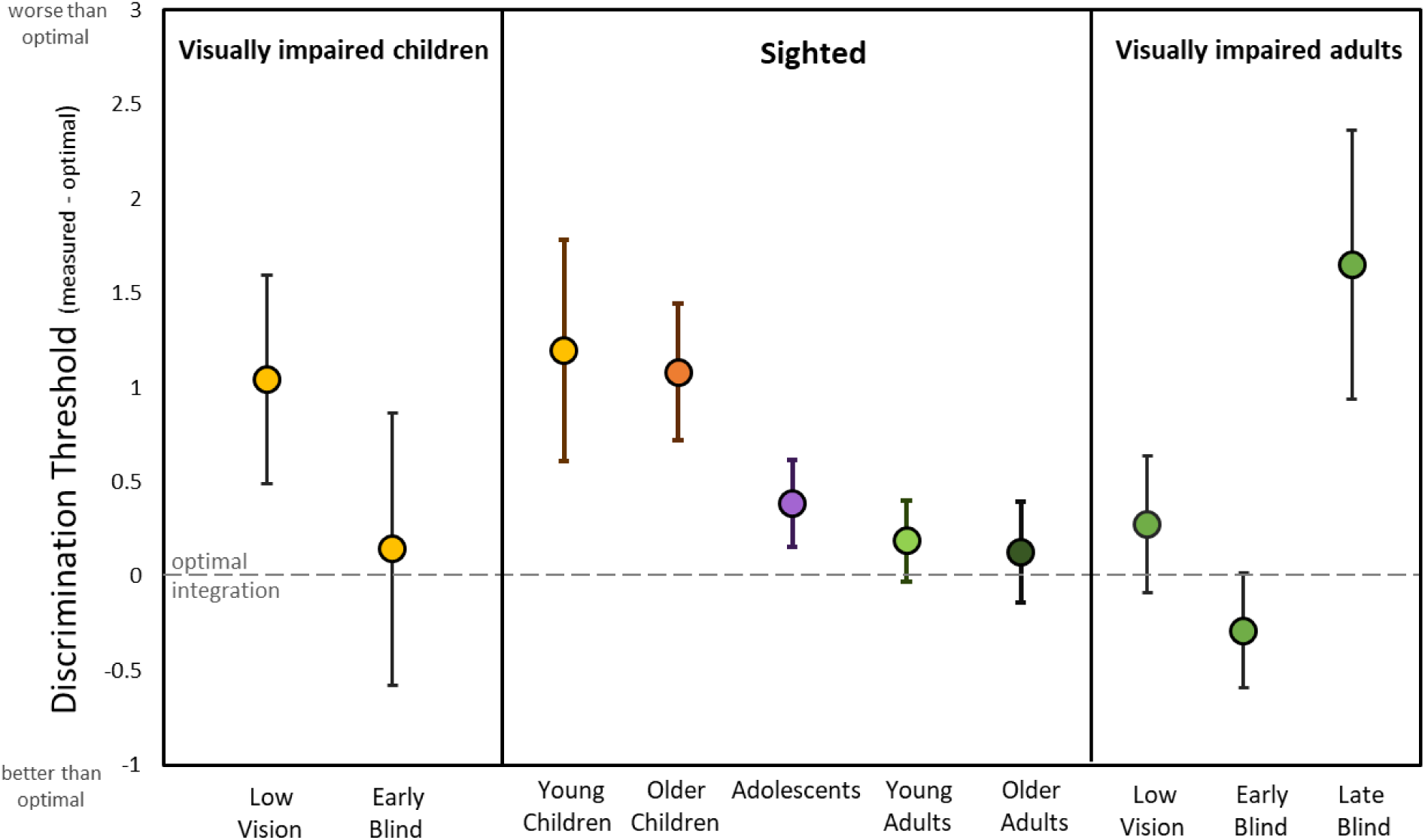
Overview figure summarizing main effects. Optimal audio-haptic integration develops across adolescence in sighted individuals. Low vision does not affect this development. Congenital and early blindness lead to an earlier development of optimal audio-haptic integration, while late blindness disrupts optimal integration.

Previous studies that reported perceptual differences between individuals with different levels of visual experience showed that congenitally blind individuals performed significantly worse than sighted individuals on different auditory and proprioceptive spatial perception tasks. At the same time, late blind and low vision individuals performed similar or even better than sighted individuals (Cappagli, Cocchi, et al., 2017; Cappagli, Finocchietti, Baud-Bovy, et al., 2017). These findings suggest that the mere presence or absence of visual input early in life affects spatial processing in the remaining senses. Interestingly, the effect of visual deprivation shows the opposite pattern in our study. A possible explanation for this opposing trend is that the present study is targeting different processes. While Cappagli et al. (2017) used a task for which vision was the most robust sense and examined the effect of visual experience on proprioception and audition separately, our study used a task for which touch was the most robust sense and we examined the effect of visual experience on the integration of touch and audition. Therefore, if vision was the most robust sense for a task, only early, but not late blindness, would affect non-visual processing later in life (Cappagli, Cocchi, et al., 2017). If touch, on the other hand, is the more robust sense for a task, early blindness should not affect non-visual processing later in life. Late blindness could, however, still affect non-visual processing if the perceptual process (e.g. non-visual multisensory integration) is dependent on the developmental consistency of sensory experience. Our results and previous findings therefore support both cross-modal compensation and cross-modal calibration. However, the results also suggest that these processes serve an adaptive purpose by allowing early sensory experience to imprint on the developing brain and preparing the developing individual for the sensory environment they are likely to experience later in life. That is, throughout the first eight years in life, the system accumulates sensory experience in order to gauge the reliability of the different sensory modalities that they will likely use later (Noppeney, Ostwald, & Werner, 2010), and to distribute modality-specific weights accordingly (Rohe, Ehlis, & Noppeney, 2019). If the early sensory environment (e.g. typical sight) does not match up with the environment that the individual experiences later in life (e.g. blindness), the system might attribute higher weights to the wrong (i.e. impaired) sensory modality.

The second aim of this study was to provide a comprehensive trajectory of the development of audio-haptic integration across the life span in sighted humans. To the best of our knowledge, only one study (Petrini et al., 2014) so far assessed how optimal audio-haptic integration develops between middle childhood (5-11 years) and young adulthood (19-35 years). They found that audio-haptic multisensory integration is not yet fully developed by the age of 11 years, with the onset of this integration remaining unknown. Our results replicate these findings, but also show that audio-haptic integration becomes more adult-like at around 13-15 years in typically sighted individuals. This is evidenced by a similar weighting of sensory cues, and a reduction in sensory uncertainty between adolescents and young adults. Arguably, the maturation of this process is still ongoing for several individuals at this age, while the majority of adolescent participants in our study benefitted from having both sensory cues available. This likely explains why the adolescent group showed a reduction of uncertainty in the audio-haptic condition compared to auditory-only or haptic-only conditions, but still differed in measured and predicted discrimination thresholds (for individual data and discussion see Supplementary material S4 and S4.2) (Jonas, Spiller, Hibbard, & Proulx, 2017; Murray, Thelen, Ionta, & Wallace, 2018; Peterzell, 2016). Finally we found that, overall, the haptic information dominated object size perception, confirming the haptic dominance for this task over other senses, which is in line with the findings of previous developmental studies (Gori et al., 2008, 2010; Petrini et al., 2014).

The summary shown in Fig. 9 suggest that the onset of adult-like integration and possibly the end of cross-modal calibration (Burr & Gori, 2012) may differ for the different senses and tasks (see also Fig. 3 in Stanley, Chen, Lewis, Maurer, & Shore, 2019). For example, the perception of temporal properties (Adams, 2016; Gori, et al., 2012) proceeds the integration of spatial characteristics (Gori et al., 2012). This is also in line with a number of studies showing that audio-visual, visuo-tactile, and audio-tactile simultaneity perception develops adult-like characteristics before the respective spatial information is integrated (Chen, Lewis, Shore, Spence, & Maurer, 2018; Chen, Shore, Lewis, & Maurer, 2016; Stanley, Chen, Lewis, Maurer, & Shore, 2019), suggesting that temporal simultaneity perception is a prequisite for the integration of spatial information. However, the onset of optimal multisensory integration also seems to depend on the sensory modality pairing that is involved in the task. For example, while audio-visual optimal integration seems to develop between 8-12 years of age (Adams, 2016; Gori et al., 2008; Gori, Sandini, et al., 2012; Nardini, Bedford, & Mareschal, 2010; Petrini et al., 2016), the integration of non-visual information does not emerge until later (Petrini et al., 2014; *present study*).

**Figure 9.**
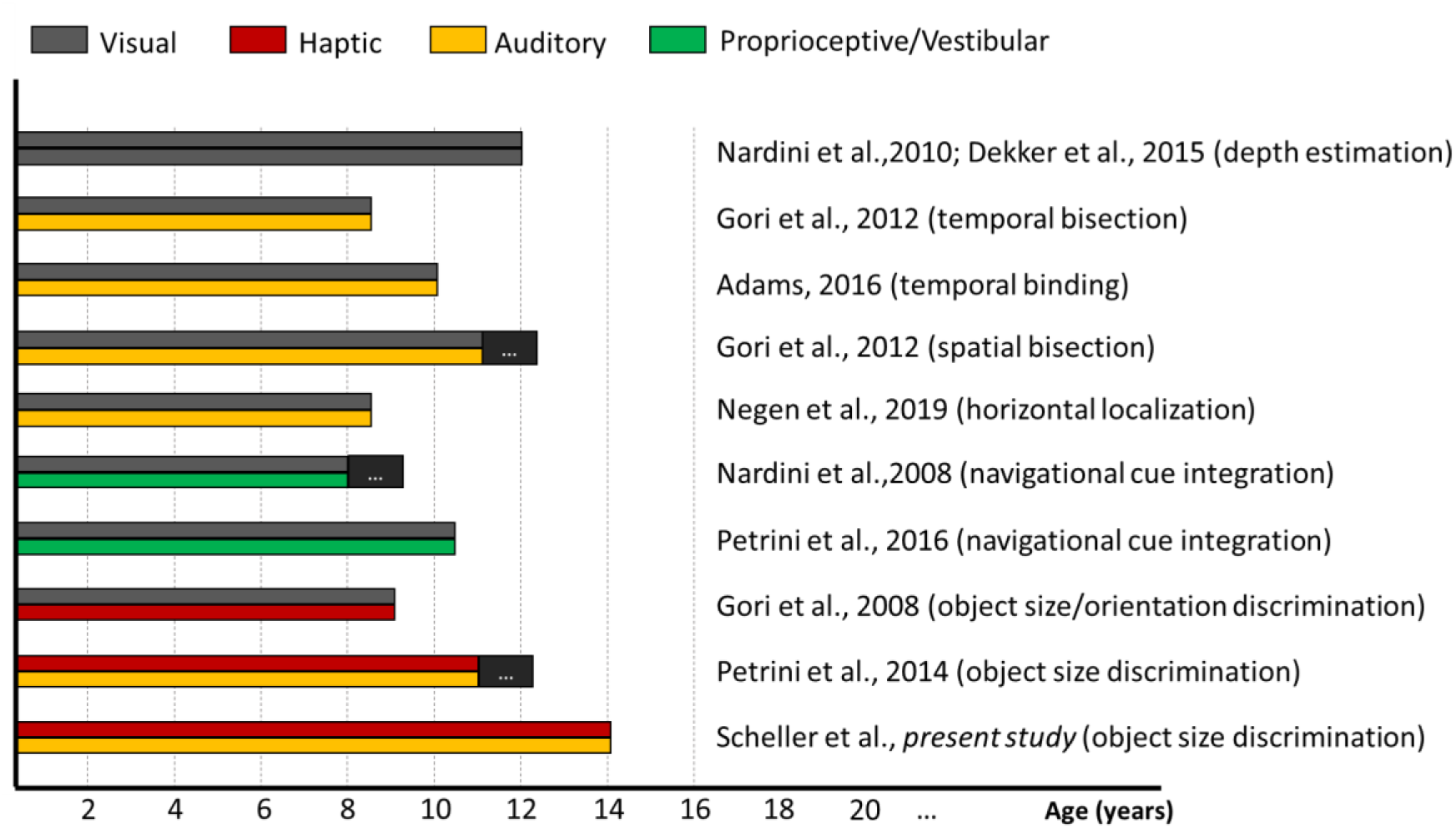
Developmental onset of adult-like multisensory integration. Reported age of onset of adult-like multisensory integration for different sensory systems. Colour combinations indicate the sensory combinations that have been tested by respective studies and tasks. All identified ages of onset fall within a period of 8-14 years, coinciding with major developments in fronto-parietal networks (Giedd et al., 1999; Gogtay et al., 2004) governing multisensory weighting (Cao et al., 2019; Rohe et al., 2019). Black boxes at the right end of the developmental trajectory indicate that multisensory integration performance has not yet reached adult-like levels at this age, but is likely to develop later (indicating the upper boundary of the age range tested in each respective study). Note that several studies did not report a concrete age of onset, but an age range during which this ability develops. The figure presents mean age of these age ranges.

The late maturation of optimal integration consistently shown by several studies (Fig. 9) could be a consequence of the late maturation of the substrates that subserve optimal multisensory integration. While early sensory processing areas mature relatively early in childhood, frontal and parietal regions have been shown to develop last, with maturational peaks around late childhood and adolescence (Giedd et al., 1999; Gogtay et al., 2004; Sowell et al., 2003). Notably, there has been long-standing evidence of the modulatory involvement of a fronto-parietal network in the optimal integration of multisensory information (Engel, Senkowski, & Schneider, 2012; Jones & Powell, 1970; Ma, Beck, Latham, & Pouget, 2006). However, specific evidence for the neural basis of multisensory reliability weighting in frontal (Cao, Summerfield, Park, Giordano, & Kayser, 2019) and parietal (Boyle, Kayser, & Kayser, 2017; Rohe et al., 2019) areas has only been provided recently. Taken together with the findings summarized in Fig. 9, this might suggest that the functional onset of optimal multisensory integration depends on the maturation of these networks, leading to a sensory-specific onset in late childhood and early adolescence. Evidence for a link between optimal cue integration within one modality and maturational changes in their processing substrate has previously been provided by Dekker and colleagues (2015).

### Conclusion

Our results show that the ability to combine audio-haptic sensory input in an optimal way does not develop before adolescence (13-17 years) in typically sighted individuals. The data further provide empirical evidence that visual experience is not necessary for non-visual optimal multisensory integration to emerge, but that consistency of sensory experience plays an important role in setting up the rules under which information is integrated later in life. They highlight that the adaptiveness of cross-modal plasticity lies in preparing the developing individual for the sensory environment they are likely to experience later in life. That is, during development, the system accumulates sensory experience in order to gauge the reliability of the different sensory modalities, and to distribute modality-specific weights accordingly. If the early sensory experience (e.g. sighted) does not match up with what the individual experiences later in life (e.g. blindness), the system might attribute higher weights to the wrong (lost or impaired) sensory modality. Our results further suggest that the calibration of the perceptual weighting system is taking place during approximately the first eight to nine years of life, highlighting the important role of early multisensory experience during this developmental period.

## Data availability statement

The data that support the findings of this study are available from the corresponding author, MS, upon reasonable request.

## Acknowledgements

We would like to thank all charities and institutions for the visually impaired that supported the study. Many thanks go especially to Jonathan Waddington from the WESC, Gemma Brimson, Denize Atan and Cathy Williams from Bristol Eye Hospital, and Emily Summers, Tania West, Rima Hussain and Konstantina Prapa from Moorfields Eye Hospital. This work would not have been possible without the dedication and participation of all visually impaired adults and children with their families. We further thank Tracey Sessions from St. John’s Catholic Primary School Bath, Izzy Hoole and Charlotte Lockyer from Clarendon Academy Trowbridge as well as all children and their parents that participated in the study. We would further like to thank Sara Garcia and Crescent Jicol for their support with setting up recruitment networks and methodological implementation. Lastly, we acknowledge the support of the National Institute for Health Research Clinical Research Network (NIHR CRN).

## Supplementary material

### S1. Participants

**Table S1.** Participant details of typically sighted adults (left) and children (right)

| Adults |  |  |  | Children |  |  |  |
| --- | --- | --- | --- | --- | --- | --- | --- |
| Participant ID | Sex | Age | Handedness | Participant ID | Sex | Age | Handedness |
| a1 | Female | 19 | right | c1 | Female | 7 | right |
| a2 | Female | 19 | right | c2 | Female | 8 | right |
| a3 | Female | 20 | right | c3 | Female | 8 | right |
| a4 | Female | 20 | right | c4 | Female | 8 | right |
| a5 | Female | 20 | left | c5 | Female | 8 | right |
| a6 | Male | 20 | right | c6 | Male | 8 | right |
| a7 | Male | 20 | right | c7 | Male | 8 | right |
| a8 | Male | 20 | left | c8 | Female | 9 | right |
| a9 | Female | 21 | right | c9 | Male | 9 | right |
| a10 | Female | 21 | right | c10 | Female | 10 | right |
| a11 | Male | 21 | right | c11 | Female | 10 | left |
| a12 | Male | 21 | right | c12 | Female | 10 | right |
| a13 | Female | 22 | left | c13 | Female | 10 | right |
| a14 | Female | 22 | right | c14 | Female | 10 | right |
| a15 | Female | 22 | right | c15 | Male | 10 | right |
| a16 | Male | 22 | right | c16 | Female | 11 | right |
| a17 | Female | 23 | right | c17 | Female | 11 | right |
| a18 | Male | 23 | right | c18 | Female | 11 | left |
| a19 | Female | 25 | right | c19 | Female | 11 | right |
| a20 | Female | 25 | right | c20 | Male | 11 | right |
| a21 | Male | 25 | left | c21 | Male | 11 | right |
| a22 | Male | 25 | right | c22 | Female | 12 | right |
| a23 | Female | 26 | right | c23 | Female | 12 | right |
| a24 | Male | 26 | right | c24 | Female | 12 | right |
| a25 | Male | 26 | right | c25 | Female | 12 | right |
| a26 | Female | 27 | right | c26 | Female | 12 | right |
| a27 | Female | 28 | right | c27 | Female | 12 | left |
| a28 | Male | 31 | right | c28 | Female | 12 | right |
| a29 | Female | 32 | right | c29 | Male | 12 | right |
| a30 | Male | 32 | right | c30 | Male | 12 | right |
| a31 | Female | 49 | right | c31 | Male | 12 | right |
| a32 | Female | 50 | right | c32 | Female | 13 | right |
| a33 | Female | 51 | right | c33 | Female | 13 | right |
| a34 | Female | 55 | right | c34 | Female | 13 | right |
| a35 | Male | 57 | right | c35 | Female | 13 | right |
| a36 | Female | 58 | right | c36 | Female | 13 | right |
| a37 | Female | 58 | right | c37 | Male | 13 | left |
| a38 | Female | 60 | right | c38 | Male | 13 | left |
| a39 | Male | 61 | right | c39 | Male | 13 | right |
| a40 | Male | 62 | right | c40 | Female | 14 | right |
| a41 | Male | 63 | left | c41 | Female | 15 | left |
| a42 | Female | 64 | right | c42 | Female | 15 | right |
| a43 | Male | 65 | right | c43 | Male | 16 | right |
| a44 | Female | 66 | right | c44 | Female | 17 | right |
| a45 | Female | 68 | right | c45 | Female | 17 | right |
| a46 | Male | 70 | right | c46 | Male | 17 | right |

Participant information of sighted adults and children can be seen in Table S1. One sighted adult participant presented symptoms of mild congenital Strabismus, however, both eyes were fully functioning and could be attended to one at a time. Therefore, despite a reduction of visual depth information through monocular vision, the participant’s visual function was unimpaired and their data were retained in the group analysis. This is further supported by research from (Kavšek & Granrud, 2012), showing that a precise object size estimation in children and adults is not dependent on the availability of binocular cues.

### S2. Methods

#### Stimuli and stimulus presentation

When creating the sound stimuli we chose to modulate sounds amplitude of a single recorded sound, instead of recording several different sounds, in order to control for the amount of information conveyed in each modality. In sound, duration, pitch or timbre might influence size discrimination strategy at different ages. Similarly, in the haptic modality, several cues (e.g. curvature, weight, height) can be used to infer object size. Hence, manipulating only one sound characteristic, amplitude, while providing differences in only one haptic property, object height, allowed for controlled provision of a similar amount of sensory information in the two modalities. Previous experiments have shown that differences in sound amplitude can be used more reliably than pitch differences for judging object size (Grassi, 2005; Petrini, Remark, Smith, & Nardini, 2014).

Stimulus presentation was controlled using Matlab with Palamedes toolbox (version 1.8.1, released: December 2, 2015; Prins & Kingdom, 2018) on a Retina MacBook Pro. A Psi adaptive staircase (Prins, 2013) was implemented for the whole stimulus range by setting up two interleaved staircases, one for the upper (49-57mm) and one for the lower side (41-49mm) of the stimulus range. Upper and lower staircase trials were randomly interleaved, leading to 15 comparisons of the standard stimulus with a larger ball, and 15 comparisons of the standard stimulus with a smaller ball, randomly presented within one block (condition). Each participant completed all four conditions. Synchronization of sound and touch was achieved using the Psychtoolbox PsychPortAudio command library (Brainard, 1997; Pelli, 1997).

#### Procedure

In all blocks, participants were instructed to attend to all the sensory information available, that is, the size they felt during the haptic-only block, the size they heard in the sound-only block, and information from both touch and hearing in the two bimodal blocks. During all patting movements, participants were not allowed to grasp or lift the ball but were instructed to keep their hands as straight and flat as possible, in order to ensure that the amount of information was limited to only one object dimension (height). At the end of each trial, they were asked to give judgements about which of the two stimuli they perceived as bigger. If unsure, they had to make a guess. The number of repetitions for each comparison stimulus depended on the participants’ previous responses and was determined by their discrimination accuracy and precision.

#### Data analysis

The two-alternative-forced-choice paradigm described in the main paper allowed us to derive discrimination thresholds from the discrimination task (Gori, Del Viva, Sandini, & Burr, 2008; Petrini et al., 2014; Rohde, van Dam, & Ernst, 2016). The discrimination threshold provides a measure of perceptual precision, as it indicates the smallest size difference that an individual can reliably detect. As higher perceptual precision leads to a reduction in noise, the discrimination threshold offers a means to quantify the reduction in sensory noise as a result of integration. Notably, when assessing optimal multisensory integration in a behaviourally beneficial sense, noise reduction is the most important assumption that needs to be met (Rohde et al., 2016).

Participants’ responses that were collected during the experiment were pre-processed in Matlab (version: R2014b, The MathWorks, USA) and further analyzed using R (version: 3.2.1). Data were sampled using an adaptive staircase (Prins & Kingdom, 2018) as this procedure allows to reliably determine psychometric characteristics while requiring fewer trials. A Psychometric function describing the probability of a participant responding that the comparison stimulus was bigger than the standard stimulus was fitted using the Quick function, which is given as

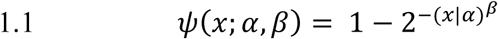

with *x* describing stimulus size, varying between the standard stimulus (49mm) and the two most extreme sizes (57mm, 41mm), *α* indicating the threshold and *β* describing the slope of the function. Both the guess and lapse rate were fixed to *γ* = 0.5 and *λ* = 0.03, respectively. Thresholds were obtained for each of the two staircases (covering the upper and the lower part of the stimulus range). They were equivalent to the point of subjective equivalence (PSE), the stimulus intensity at which a participant cannot tell the difference between two stimuli (i.e. 50% of responses “bigger”). This point can further be used to calculate the Just Noticeable Difference (JND) for the overall psychometric functions (see Fischer & Whitney, 2014). The just noticeable difference indicates the minimum size difference that can reliably be discriminated, and can be extracted as:

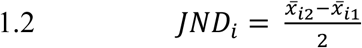

Where *x̅_i1_* denotes the absolute detection threshold (PSE) for the lower side of the stimulus range and *x̅_i2_* for the upper side for each experimental condition *i*. The two PSEs are the points at which 25% and 75% of the comparison stimuli were rated as “bigger”, respectively. Both JND and PSE for each participant and condition were exported for further processing in R. Discrimination thresholds (σ) were calculated based on each individual JND as:

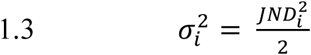

Maximum Likelihood Estimation was used to calculate predicted bimodal precision according to optimal integration, as given by:

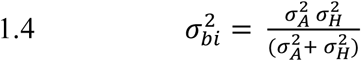

With 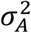 indicating the measured variance in the auditory performance and 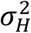 indicating the measured variance in the haptic performance.

In order to assess the developmental trajectory of optimal multisensory integration by means of optimal sensory noise reduction, we compared the measured discrimination thresholds of children and adolescents, as well as older adults, with a group of younger adults (18-44 year old). This is because optimal multisensory integration has been commonly established in this particular demographic. In order to quantify the multisensory benefit in terms of optimal noise reduction for each age group, we computed the difference between the measured bimodal discrimination threshold and the discrimination threshold predicted by MLE (Δ_measured-predicted_) for each individual separately. This measure provides a quantified estimation of the perceptual benefit that is gained through multisensory processes alone, as each individual’s MLE prediction is calculated based on their unisensory precision for touch and hearing. It thereby takes inter-individual variation in the precision of the two sensory systems into account. The lower Δ_measured-predicted_ is for each individual, the more they benefit from noise reduction through multisensory integration. After assessing the development of multisensory benefit in the sighted population, we compared adults and children with low vision or blindness with different ages of onset to the respective age groups.

Weights attributed to the haptic cue (*ω_H_*) were calculated from discrimination thresholds in the congruent condition via:

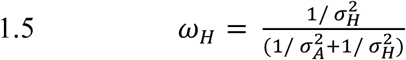

Please note that when auditory and haptic cues are presented simultaneously (as in the bimodal conditions) the auditory weight can be calculated as 1 - *ω_H_*.

Furthermore, shifts in PSEs were assessed for the incongruent condition in which a haptic-auditory conflict was introduced in order to assess whether biases in sensory cue selection change across development. Here, weights derived from PSEs were calculated from:

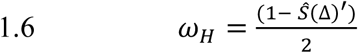

With *Ŝ*(Δ)^′^ indicating the slope of a linear regression of PSEs for all values of Δ. For more information see Supplemental Data in (Gori et al., 2008).

### S3. Results – Test assumptions

#### Discrimination thresholds

##### Sighted participants

With the exception of the youngest children group for sound discrimination thresholds (p = 0.01), the thresholds from all age groups and in all conditions was normally distributed (p > .05) as assessed by a Shapiro-Wilk test. Levene’s test for homogeneity of variances indicated that variances were not different across age groups (p > .05). No data points lying outside 1.5 IQR were detected, assuming no outliers. As analysis of variance is robust to violations of normality, we conducted a mixed factorial analysis of variance with the conditions as within-subjects factor and age group as between-subjects factor.

##### Visually impaired participants

Due to the small sample sizes for all visually impaired adult groups (all n = 3) we used non-parametric Mann-Whitney U-tests to compare visually impaired adults with sighted adults. For comparability, we used the same test to compare low vision children with sighted children. As the blind children (n = 2) and blind adolescent groups (n = 2) were even smaller in size, we conducted single-case comparisons of these individuals with the respective age-matched sighted children and adolescent groups using a Crawford-Howell t-test for single case-control comparisons (Crawford, Garthwaite, & Porter, 2010).

#### Multisensory benefit (Δ_measured-predicted_)

##### Sighted participants

In order to compare between age groups, parametric test assumptions were assessed for Δ_measured-predicted_. We identified one outlier in the young adult group with a Δ_measured-predicted_ outside of 1.5 IQR from the upper quartile of the distribution, which was due to an exceptionally low threshold in the haptic condition. After removing the outlier, all other assumptions of parametric testing were met in all sighted age groups. We therefore conducted independent, Bonferroni-corrected t-tests to assess the differences in multisensory integration between young adults and other developmental age groups.

##### Visually impaired participants

We used non-parametric Mann-Whitney U-tests and Crawford-Howell case-control comparison t-tests for comparing visually impaired with sighted age-matched groups as described above for the discrimination thresholds.

#### Incongruent condition

##### Sighted participants

Shapiro-Wilk tests indicated that, with the exception of sighted young adults (p = 0.02), the data from all age groups was normally distributed (p > .05). Levene’s test for homogeneity of variances indicated that variances were not different across age groups (p > .05). We identified three outliers (>1.5 IQR) in the age groups 10-12 years, 13-17years, and 45-70 years in the sighted sample, indicating higher thresholds than the rest of the group. However, as individual differences in the response to incongruent stimulus pairings are meaningful in that they indicate different sensory combination strategies (i.e. integration or switching between modalities), these values were retained in the analysis.

##### Visually impaired participants

We used non-parametric Mann-Whitney U-tests to compare sensory weights derived from discrimination thresholds and from PSEs.

### S4. Results – Individual performance and integration strategies

In order to examine how individuals combined audio and haptic cues in the bimodal condition, we plotted ratios of single-cue variances (auditory/haptic) against ratios of bimodal-to haptic-cue variances for the different age groups (see Figure S8.A). The red and green line indicate predictions for either relying mostly on the worse (red) or on the best (green) sensory cue. Most individual data of adults and 13-17year olds falls below the green line, with group averages decreasing on the ordinate, indicating that they benefitted from combining sensory cues in the bimodal condition. Children in both age groups, 7-9years and 10-12years, on the other hand, show a bimodal-to haptic-variance ratio that can be approximated by using the worse sensory cue. Overall, sound cues were more reliably used in the older age groups, despite haptic information remaining the more reliable cue. This indicated an improvement in reliability and use of auditory information in the bimodal condition with age. Results for the two younger age groups and the young adults reliably replicate key findings of an earlier study using a similar paradigm (Petrini et al., 2014).

**Figure S8.1.**
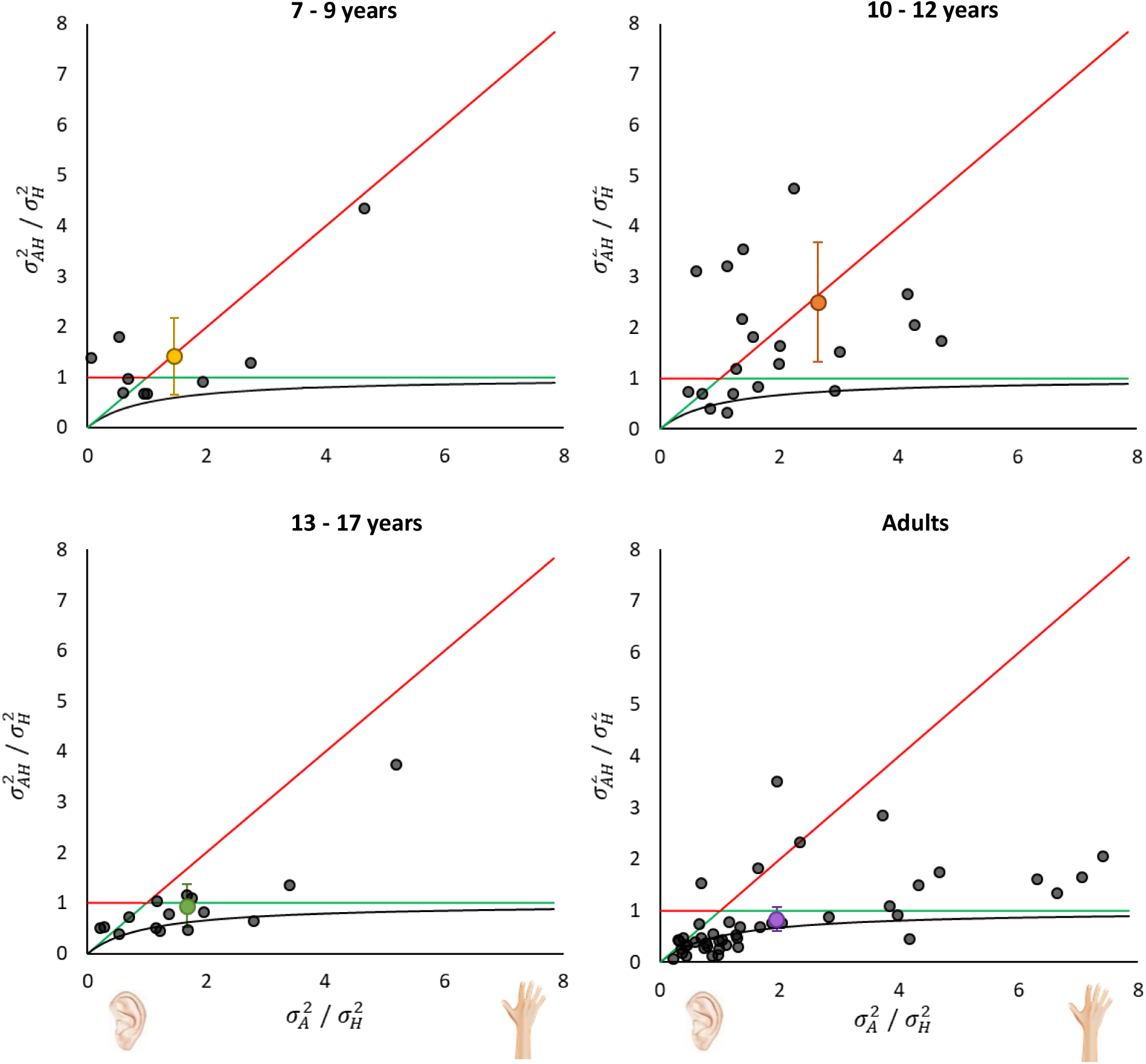
Unimodal and bimodal variance ratios of sighted individuals. Individual data (black circles) and group averages (colored circles) of variance ratios for auditory and haptic single-cues (σA/σH) and bimodal-to-haptic cues (σAH/σH). Error bars indicate 95% CIs. Higher ratios along the abscissa indicate lower variance for touch, meaning that touch is more reliable than hearing. Lower ratios along the y-axis indicate an improvement with both cues compared to touch alone. For comparison, model predictions are plotted based on using the single worst cue (red line), the single best cue (green line), or the integration of both cues following the Bayesian model (black line).

Single-cue variances (auditory/haptic) were further plotted against ratios of bimodal-to haptic-cue variances for the visually impaired individuals, as depicted in Figure S4.. In the group of adults that lost their sight after eight years of life (late blind), integration of audio and haptic cues did not, on average, lead to an improvement in performance. They did not benefit from having a second cue available. Contrarily, the majority of congenitally and early blind individuals, who lost their vision within the first eight years of life, showed a perceptual benefit in having multiple sensory cues available by means of a decrease in discrimination threshold (higher precision) in the bimodal condition compared to the more reliable unisensory condition. This improvement was evident for both the adults and children in this group, with the exception of one child – indicated as a black triangle in Figure S4.. This 13-year old individual showed a high reliance on touch compared to audition. During testing, we observed this individual to employ different approaches of using active touch between the bimodal and haptic condition, which was likely due to experience. That is, in the bimodal condition (which was also presented as the first block to them), they tapped the ball rapidly, similar to a button press, in order to elicit the sound, and judged object size based on sound, without paying much attention to touch. This was also verbally reported by the individual after the testing session. Contrarily, in the haptic condition, which they completed last, this individual repeatedly grasped the ball slowly, thereby gaining more information about object size from touch. This might explain why this individual did not gain much perceptual precision in the bimodal compared to the haptic only condition (see Figure S4.). Due to the limited sample size of congenitally and early blind children (n = 4), we decided to include this individual’s responses in the analysis. However, it should be noted that this individual might have used haptic input differently across the conditions, and that performance is likely better in early blind children than the group average suggests.

**Figure S4.2.**
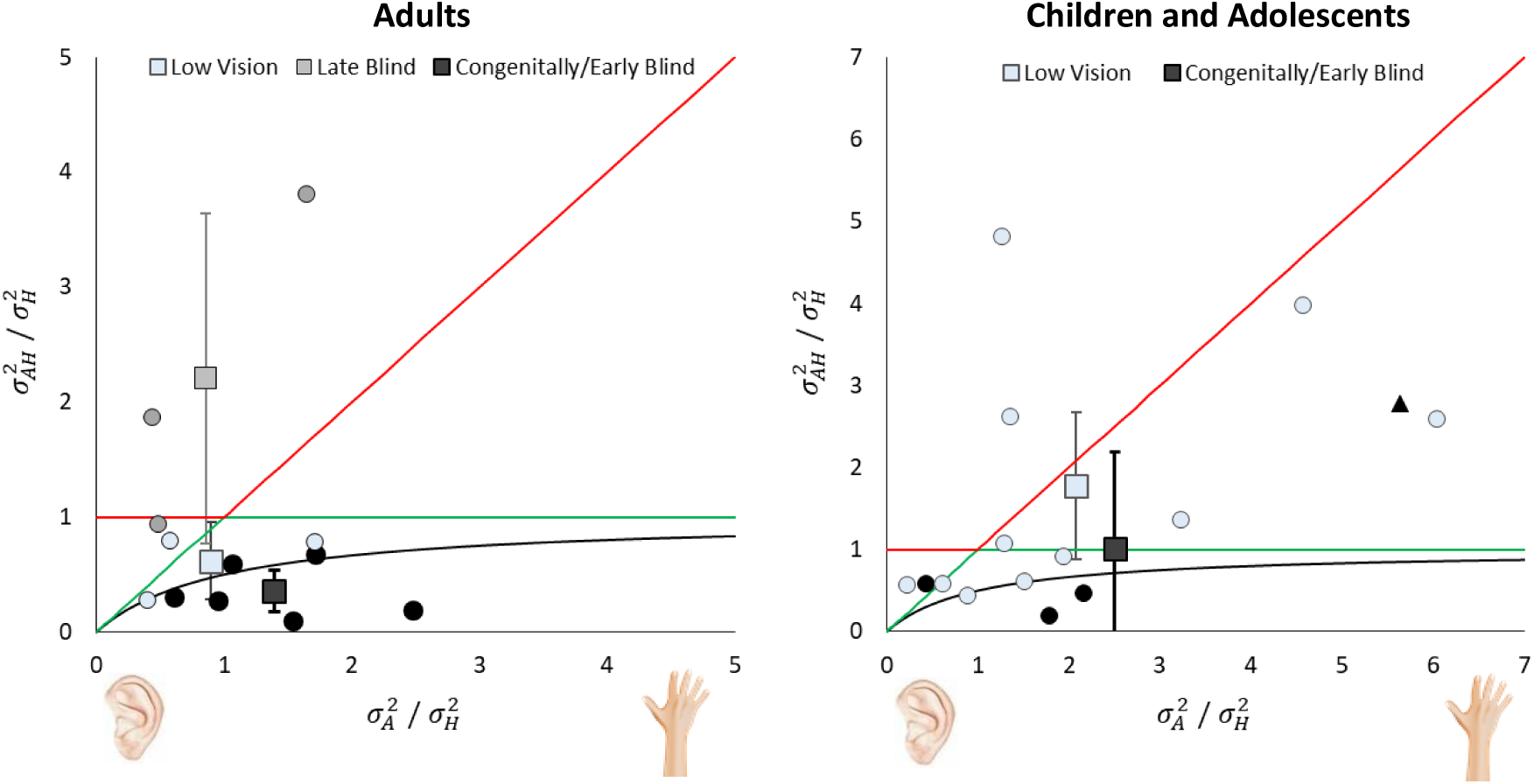
Unimodal and bimodal variance ratios of visually impaired individuals. Variance ratios for auditory and haptic single-cues (σA/σH) and bimodal-to-haptic cues (σAH/σH) for visually impaired adults (left) and children (right) with different levels of visual experience. Higher ratios along the abscissa indicate lower variance for touch, meaning that touch is more reliable than hearing. Lower ratios along the y-axis indicate an improvement with both cues compared to touch alone. Squares indicate group average with error bars representing 95% CIs. Model predictions are plotted based on using the single worst cue (red line), the single best cue (green line), or the integration of both cues following the Bayesian model (black line). Black triangle in right panel marks blind adolescent individual that used active touch differently (see main text above).

#### S4.2. Results – Typically sighted individuals - Adolescent groups

Due to the small sample size of the older adolescent group (*n* = 4), both groups, the younger adolescents (13-15 years) and older adolescents (16-17years) were combined in the main analysis. However, as depicted in Figure S4.2 we can observe a clear difference in the bimodal thresholds versus predicted thresholds between the age groups of 10-12 years and 13-15 years. The younger and older adolescent groups show similar bimodal discrimination thresholds, justifying the grouping of younger and older adolescents into one group (13-17 years). To confirm the developmental effect reported in our main analysis, we carried out an additional t-test between young adults and young adolescents (13-15 year olds). This test indicated no significant difference in Δ_measured-predicted_ (t(22) = 1.63, p =.292, d_unb_ = 0.383), suggesting the developmental onset of adult-like MSI at 13-15 years of age.

Figure S4.2 shows the relationship between measured bimodal discrimination threshold and discrimination threshold predicted by MLE for typically sighted individuals, with adolescent groups split into younger (13-15 year old) and older (16-17 year old) adolescents.

**Figure S4.2.**
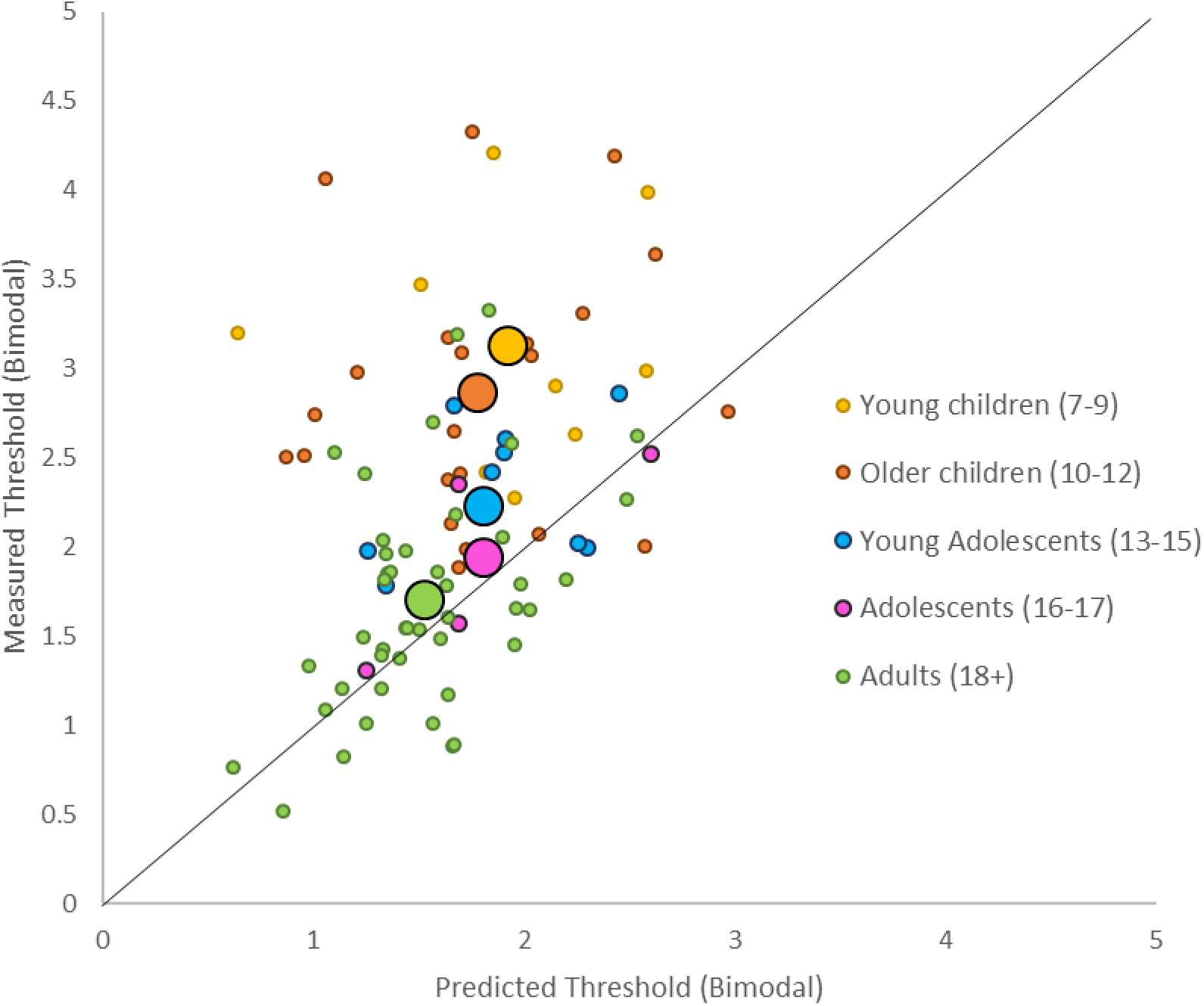
Predictability of measured bimodal discrimination thresholds by MLE. Measured bimodal thresholds for all individuals in five different age groups plotted against MLE-predicted threshold. Small circles represent individual data, while large circles indicate group means. Data points falling closer to the black, diagonal line indicate observed bimodal thresholds being more similar to MLE prediction. Adolescent groups are separated into younger and older adolescents to allow a clearer observation of the developmental trend. Due to the overlap, and to aid data visualization, older and younger adults have been combined into one group.

On the individual level, the youngest age at which we found children to optimally integrate audio-haptic information was 10 years. However, the large majority of 10-12-year-old children did not integrate both cues to reduce sensory uncertainty. This also highlights the individual differences in the onset of sensory uncertainty reduction, which likely depend on factors such as early sensory experience and cognitive maturation (see Nardini, Begus, & Mareschal, 2013; Petrini et al., 2014). As most studies assessing the extent of optimal multisensory integration quantitatively focus on averaged group measures, individual differences are often ignored while they can provide useful information about potential mechanisms that lead to differences in developmental onset.

### S5. Results – Sensory weights

To assess how individuals weighted sensory information, and whether cue combination strategies differed between the different developmental groups, we compared individuals’ performances between the bimodal congruent and incongruent condition. This allows us to disentangle whether participants weighted haptic or auditory information more strongly in order to reduce uncertainty in the bimodal congruent condition, and whether they relied more on auditory or haptic information when the cues gave conflicting information.

#### S5.1. Typically sighted individuals

Mean weights derived from thresholds (see figure S5.1, left panel) were not significantly different from 0.5 for all age groups (7-9: *t*(8) = 2.31, *p* = 0.868; 13-17: *t*(14) = 2.14, *p* = 0.326; 18-44: *t*(29) = 2.05, *p* = 0.108; 45-70: *t*(15) = 2.13, *p* = 0.586), with exception from the 10-12 year olds. The latter showed significantly higher haptic weighting during bimodal integration (*t*(21) = 2.08, *p* = 0.001). Mean weights derived from PSEs in the incongruent condition (see figure S5.1 right panel) indicate a higher weighting of haptic information for all age groups (7-9: *t*(8) = 3.54, *p* = .008 ; 10-12: *t*(21) = 2.13 *p* = .045; 13-17: *t*(14) = 3.53, *p* = .003; 18-44: *t(29*) = 5.82, *p* < .001), except for the older adults (*t*(15) = 0.07, *p* = .943). This is in line with the findings from Petrini and colleagues (2014), showing no difference in sensory weighting between children and young adults.

**Figure S5.1.**
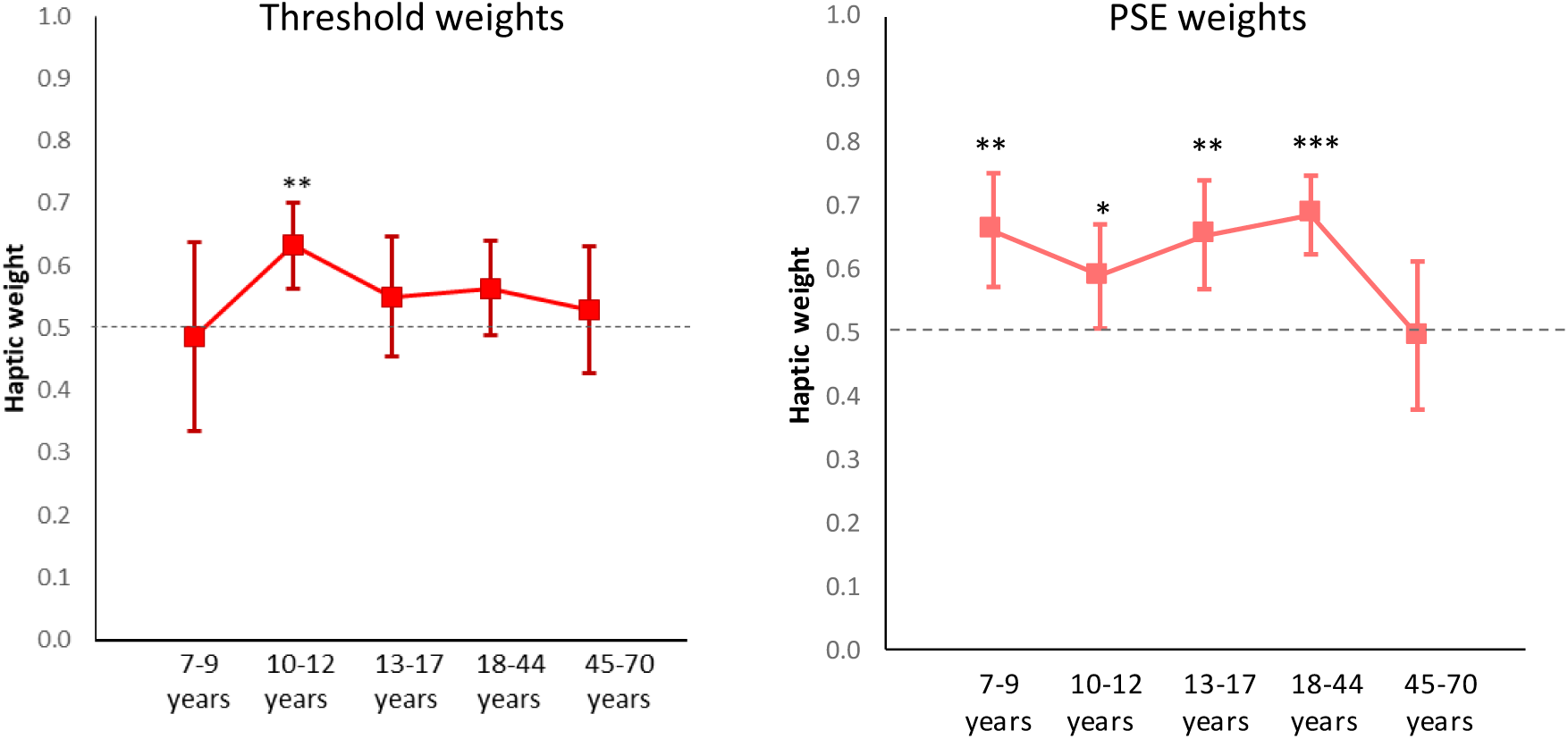
Haptic weights in sighted individuals. Mean haptic weights for the different developmental age groups derived from discrimination thresholds in the congruent condition (left) and PSE shifts in the incongruent condition (right). Values above the dashed line at y =0.5 indicate haptic dominance, while values below this line indicate auditory dominance. Figure shows the mean weights for each age group with error bars indicating 95% CI. * = p < .05; ** = *p* < .01; *** = *p* < .001.

#### S5.2. Visually impaired and blind individuals

Due to small group sizes we show the individual weights derived from thresholds in figure S5.2, upper two panels) and derived from PSE shifts (lower two panels) for individuals with low vision and for blind individuals. Mean threshold weights were not significantly different from 0.5 in any of the groups (*p* > .05), indicating that neither haptic nor auditory modalities dominated significantly. However, the figures show a similar trend to sighted individuals in both low vision and blind individuals, with children and young adults weighting the haptic cue more strongly while older adults weight the auditory cue more, independently of visual experience.

**Figure S5.2.**
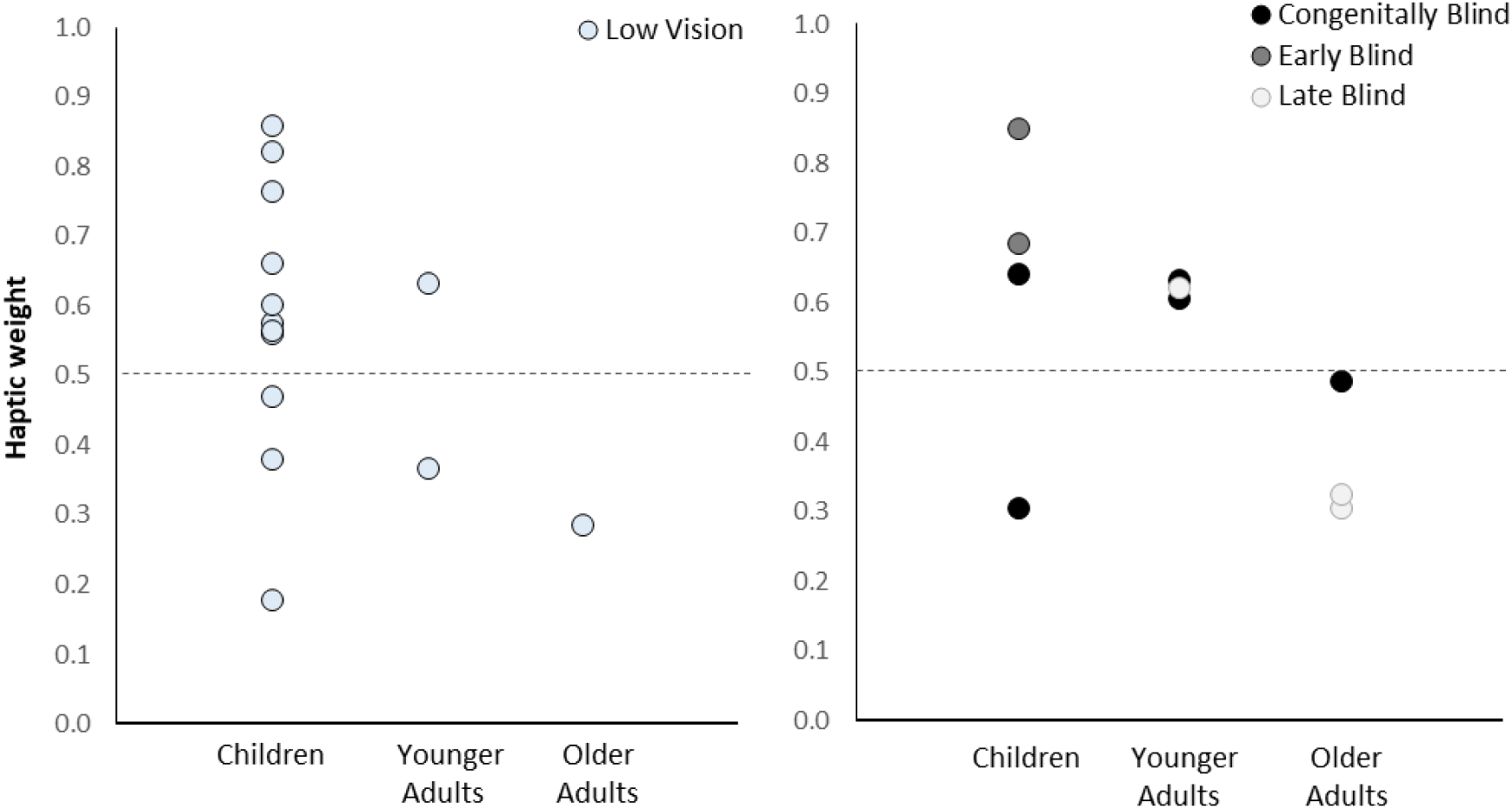
Haptic weights in visually impaired individuals. Individual haptic weights for the different age and vision groups derived from discrimination thresholds in the congruent condition (upper panels) and PSE shifts in the incongruent condition (lower panels). Values above the dashed line at y =0.5 indicate haptic dominance, while values below this line indicate auditory dominance.

### S6. Results – Bimodal congruency

#### S6.1. Typically Sighted individuals

To assess whether differences in size discrimination thresholds for congruent and incongruent conditions differed between the age groups, we carried out a mixed factorial ANOVA, using condition as within-subjects factor and age group as between-subjects factor. This revealed a significant main effect of age (*F*(4,87) = 14.64, *p* < .001), as well as a significant interaction between age and condition (*F*(8,87) = 5.67, *p* < .001; see Figure S6.1). Follow-up, Bonferroni-corrected t-tests indicated that younger adults and older adults showed significantly lower thresholds in the congruent condition, compared to the incongruent condition (18-44years: *t*(29) = 3.72, *p* = .004, *d_unb_* = 0.67; 45-70years: *t*(15) = 3.67, *p* = .01, *d_unb_* = .895). This was not the case for the two children groups (7-9 years: *t*(8) = 1.85, *p* = .504, *d_unb_* =0.589; 10-12 years: *t*(21) = 0.99, *p* = 1, *d_unb_* = 0.021), nor for the adolescent group (*t*(14) = 0.07, *p* = 1, *d_unb_* = 0.017). This supports the findings that adults increased perceptual precision by integrating congruent information and engaged in strategy switching more frequently in the incongruent condition. Children, on the other hand, were more likely to base their size discrimination judgement on one sense rather than combining both senses.

**Figure S6.1.**
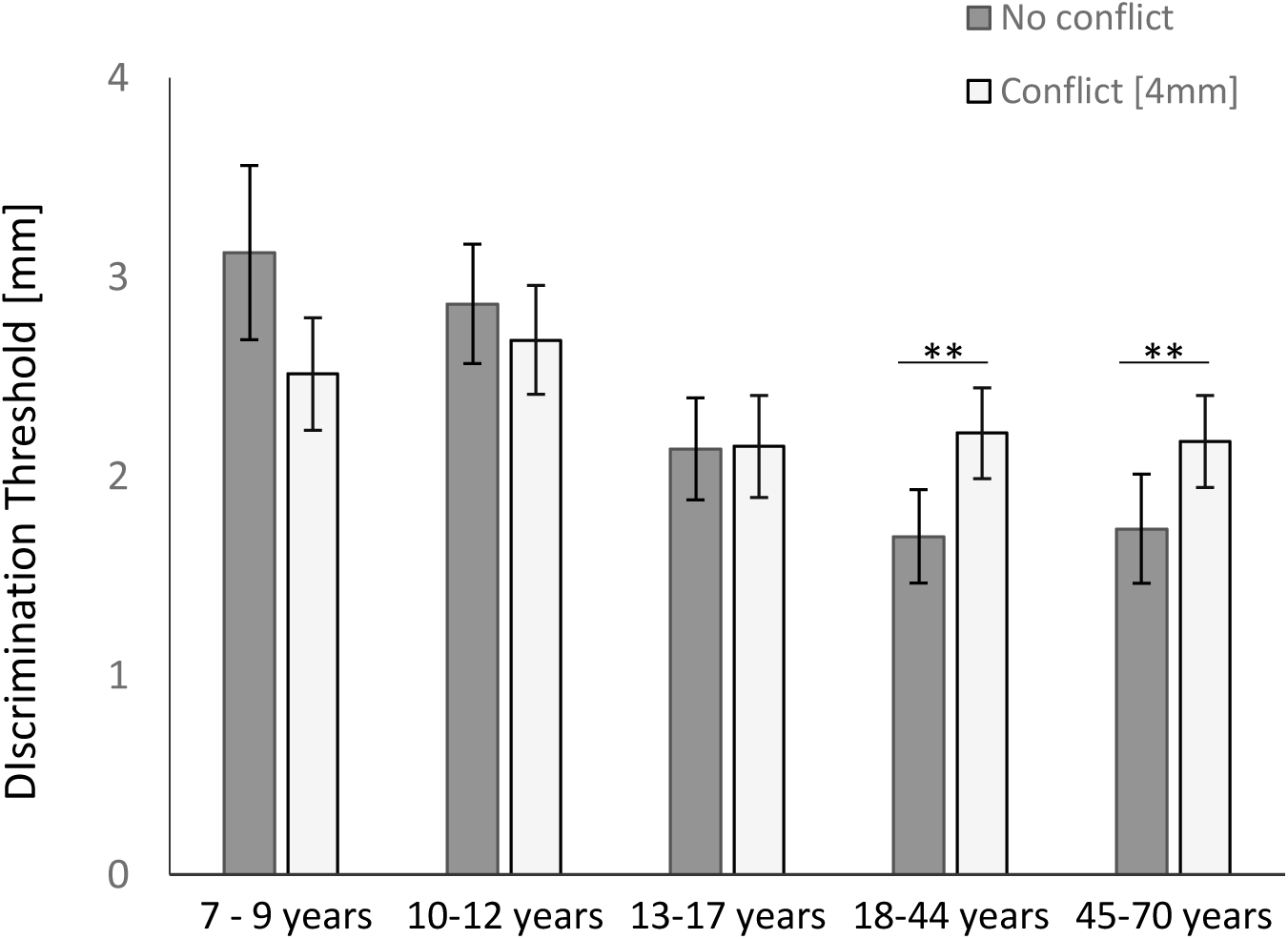
Effect of congruency on size discrimination performance in sighted individuals. Mean discrimination thresholds for the different age groups in the bimodal congruent (dark grey bar) and the bimodal incongruent (light grey bar) condition. Error bars indicate 95% CI. ** = p ≤ 0.01.

#### S6.2. Visually impaired and blind individuals

To assess whether differences in size discrimination thresholds for congruent and incongruent conditions differed between the different vision groups, we carried out non-parametric Mann-Whitney U-tests between in three different groups: low vision individuals, blind individuals that integrated optimally (congenitally blind and early blind adults and children), and blind individuals that did not integrate optimally (late blind adults). We grouped the early and congenitally blind individuals in order to allow us to increase power, and because congenitally and early blind children and adults use audio-haptic information very similarly. Please note that Figure S6.2 shows all six separate vision groups.

The tests revealed no difference between congruent and incongruent condition for the low vision group (*U* = 65, *p* = .191) nor for the late blind group (*U* = 3, *p* = .50). The early blind group showed a similar trend to sighted adults, with lower discrimination thresholds in the congruent compared to incongruent condition (*U* = 4, *p* = .014), indicating that precision was higher when individuals integrated sensory cues compared to task switching or focusing on only one sense at a time.

**Figure S6.2.**
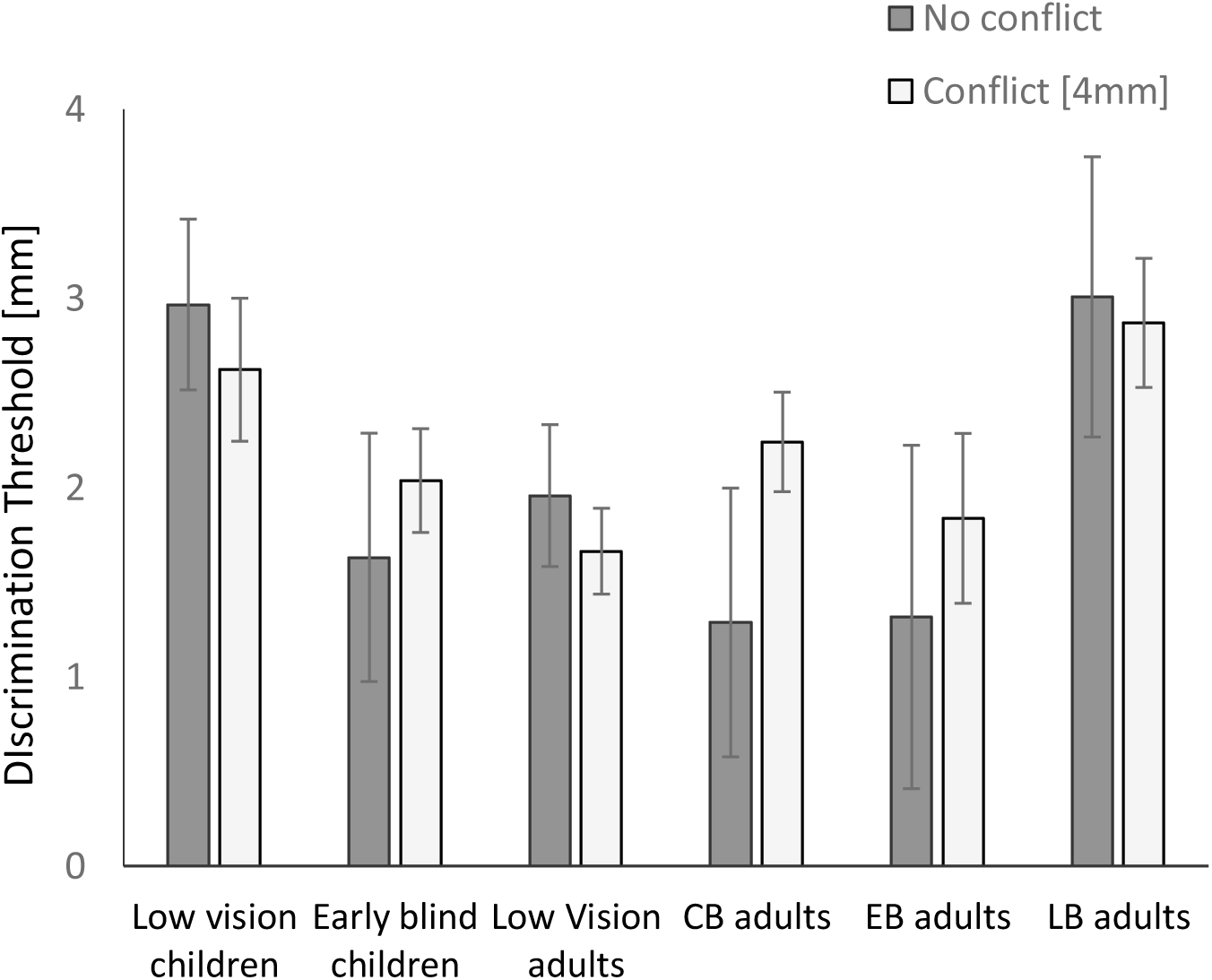
Effect of congruency on size discrimination performance in visually impaired individuals. Mean discrimination thresholds for the different vision groups in the bimodal congruent (dark grey bar) and the bimodal incongruent (light grey bar) condition. Error bars indicate 95% CI.

## References

Adams, W. J. (2016). The development of audio-visual integration for temporal judgements. PLoS Computational Biology, 12(4), e1004865. https://doi.org/10.1371/journal.pcbi.1004865

Amedi, A., Raz, N., Pianka, P., Malach, R., & Zohary, E. (2003). Early “visual” cortex activation correlates with superior verbal memory performance in the blind. Nature Neuroscience, 6(7), 758–766. https://doi.org/10.1038/nn1072

Bedny, M., Pascual-Leone, A., Dravida, S., & Saxe, R. (2012). A sensitive period for language in the visual cortex: Distinct patterns of plasticity in congenitally versus late blind adults. Brain and Language, 122(3), 162–170. https://doi.org/10.1016/j.bandl.2011.10.005

Ben Porquis, L., Finocchietti, S., Zini, G., Cappagli, G., Gori, M., & Baud-Bovy, G. (2017). ABBI: A wearable device for improving spatial cognition in visually-impaired children. In 2017 IEEE Biomedical Circuits and Systems Conference (BioCAS) (pp. 1–4). IEEE. https://doi.org/10.1109/BIOCAS.2017.8325128

Boersma, P. (2001). PRAAT, a system for doing phonetics by computer. Glot International, 5(9/10), 341–345. https://doi.org/10.1097/AUD.0b013e31821473f7

Boyle, S. C., Kayser, S. J., & Kayser, C. (2017). Neural correlates of multisensory reliability and perceptual weights emerge at early latencies during audio-visual integration. European Journal of Neuroscience, 46(10), 2565–2577. https://doi.org/10.1111/ejn.13724

Burr, D., & Gori, M. (2012). Multisensory Integration Develops Late in Humans. In M. M. Murray & M. T. Wallace (Eds.), The Neural Bases of Multisensory Processes (pp. 345–362). Boca Raton (FL): CRC Press/Taylor & Francis.

Cao, Y., Summerfield, C., Park, H., Giordano, B. L., & Kayser, C. (2019). Causal Inference in the Multisensory Brain. Neuron, 102(5), 1076–1087.e8. https://doi.org/10.1016/j.neuron.2019.03.043

Cappagli, G., Cocchi, E., & Gori, M. (2017). Auditory and proprioceptive spatial impairments in blind children and adults. Developmental Science, 20(3). https://doi.org/10.1111/desc.12374

Cappagli, G., Finocchietti, S., Baud-Bovy, G., Cocchi, E., & Gori, M. (2017). Multisensory Rehabilitation Training Improves Spatial Perception in Totally but Not Partially Visually Deprived Children. Frontiers in Integrative Neuroscience, 11, 29. https://doi.org/10.3389/fnint.2017.00029

Cappagli, G., Finocchietti, S., Cocchi, E., & Gori, M. (2017). The Impact of Early Visual Deprivation on Spatial Hearing: A Comparison between Totally and Partially Visually Deprived Children. Frontiers in Psychology, 8, 467. https://doi.org/10.3389/fpsyg.2017.00467

Champoux, F., Collignon, O., Bacon, B. A., Lepore, F., Zatorre, R. J., & Théoret, H. (2011). Early- and Late-Onset Blindness Both Curb Audiotactile Integration on the Parchment-Skin Illusion. Psychological Science, 22(1), 19–25. https://doi.org/10.1177/0956797610391099

Chen, Y.-C., Lewis, T. L., Shore, D. I., Spence, C., & Maurer, D. (2018). Developmental changes in the perception of visuotactile simultaneity. Journal of Experimental Child Psychology, 173, 304–317. https://doi.org/10.1016/j.jecp.2018.04.014

Chen, Y.-C., Shore, D. I., Lewis, T. L., & Maurer, D. (2016). The development of the perception of audiovisual simultaneity. Journal of Experimental Child Psychology, 146, 17–33. https://doi.org/10.1016/j.jecp.2016.01.010

Collignon, O., Dormal, G., Albouy, G., Vandewalle, G., Voss, P., Phillips, C., & Lepore, F. (2013). Impact of blindness onset on the functional organization and the connectivity of the occipital cortex. Brain, 136(9), 2769–2783. https://doi.org/10.1093/brain/awt176

Collignon, O., Dormal, G., De Heering, A., Lepore, F., Lewis, T. L., & Maurer, D. (2015). Long-Lasting Crossmodal Cortical Reorganization Triggered by Brief Postnatal Visual Deprivation. Current Biology, 25(18), 2379–2383. https://doi.org/10.1016/j.cub.2015.07.036

Coluccia, E., Mammarella, I. C., & Cornoldi, C. (2009). Centred egocentric, decentred egocentric, and allocentric spatial representations in the peripersonal space of congenital total blindness. Perception, 38(5), 679–693. https://doi.org/10.1068/p5942

Crawford, J. R., Garthwaite, P. H., & Porter, S. (2010). Point and interval estimates of effect sizes for the case-controls design in neuropsychology: Rationale, methods, implementations, and proposed reporting standards. Cognitive Neuropsychology, 27(3), 245–260. https://doi.org/10.1080/02643294.2010.513967

Cumming, G. (2012). Understanding the new statistics: Effect sizes, confidence intervals, and meta-analysis. New York, NY: Routledge.

de Klerk, C. C. J. M., Johnson, M. H., Heyes, C. M., & Southgate, V. (2015). Baby steps: Investigating the development of perceptual-motor couplings in infancy. Developmental Science, 18(2), 270–280. https://doi.org/10.1111/desc.12226

Engel, A. K., Senkowski, D., & Schneider, T. R. (2012). Multisensory Integration through Neural Coherence. The Neural Bases of Multisensory Processes. CRC Press/Taylor & Francis.

Ernst, M. O., & Banks, M. S. (2002). Humans integrate visual and haptic information in a statistically optimal fashion. Nature, 415(6870), 429–433. https://doi.org/10.1038/415429a

Garcia, S., Petrini, K., Rubin, G. S., Da Cruz, L., & Nardini, M. (2015). Visual and non-visual navigation in blind patients with a retinal prosthesis. PLoS ONE, 10(7), e0134369. https://doi.org/10.1371/journal.pone.0134369

Geldart, S., Mondloch, C. J., Maurer, D., De Schonen, S., & Brent, H. P. (2002). The effect of early visual deprivation on the development of face processing. Developmental Science, 5, 490–501.

Giedd, J. N., Blumenthal, J., Jeffries, N. O., Castellanos, F. X., Liu, H., Zijdenbos, A., … Rapoport, J. L.. (1999). Brain development during childhood and adolescence: a longitudinal MRI study. Nature Neuroscience, 2(10), 861–863. https://doi.org/10.1038/13158

Gogtay, N., Giedd, J. N., Lusk, L., Hayashi, K. M., Greenstein, D., Vaituzis, A. C., … Thompson, P. M.. (2004). Dynamic mapping of human cortical development during childhood through early adulthood. Proceedings of the National Academy of Sciences of the United States of America, 101(21), 8174–8179. https://doi.org/10.1073/pnas.0402680101

Goldreich, D., & Kanics, I. M. (2003). Tactile acuity is enhanced in blindness. The Journal of Neuroscience, 23(8), 3439–3445.

Gori, M., Cappagli, G., Tonelli, A., Baud-Bovy, G., & Finocchietti, S. (2016). Devices for visually impaired people: High technological devices with low user acceptance and no adaptability for children. Neuroscience & Biobehavioral Reviews, 69, 79–88. https://doi.org/10.1016/J.NEUBIOREV.2016.06.043

Gori, M., Del Viva, M., Sandini, G., & Burr, D. C. (2008). Young Children Do Not Integrate Visual and Haptic Form Information. Current Biology, 18(9), 694–698. https://doi.org/10.1016/j.cub.2008.04.036

Gori, M., Sandini, G., & Burr, D. (2012). Development of Visuo-Auditory Integration in Space and Time. Frontiers in Integrative Neuroscience, 6(September), 77. https://doi.org/10.3389/fnint.2012.00077

Gori, M., Sandini, G., Martinoli, C., & Burr, D. (2010). Poor Haptic Orientation Discrimination in Nonsighted Children May Reflect Disruption of Cross-Sensory Calibration. Current Biology, 20(3), 223–225. https://doi.org/10.1016/j.cub.2009.11.069

Gori, M., Sandini, G., Martinoli, C., & Burr, D. C. (2014). Impairment of auditory spatial localization in congenitally blind human subjects. Brain, 137(1), 288–293. https://doi.org/10.1093/brain/awt311

Gori, M., Tinelli, F., Sandini, G., Cioni, G., & Burr, D. (2012). Impaired visual size-discrimination in children with movement disorders. Neuropsychologia, 50(8), 1838–1843. https://doi.org/10.1016/j.neuropsychologia.2012.04.009

Gougoux, F., Lepore, F., Lassonde, M., Voss, P., Zatorre, R. J., & Belin, P. (2004a). Neuropsychology: Pitch discrimination in the early blind. Nature, 430(6997), 309–309. https://doi.org/10.1038/430309a

Gougoux, F., Lepore, F., Lassonde, M., Voss, P., Zatorre, R. J., & Belin, P. (2004b). Pitch discrimination in the early blind. Nature, 430(6997), 309–309. https://doi.org/10.1038/430309a

Gougoux, F., Zatorre, R. J., Lassonde, M., Voss, P., & Lepore, F. (2005). A Functional Neuroimaging Study of Sound Localization: Visual Cortex Activity Predicts Performance in Early-Blind Individuals. PLoS Biology, 3(2), e27. https://doi.org/10.1371/journal.pbio.0030027

Guerreiro, M. J. S., Putzar, L., & Röder, B. (2016). The Effect of Early Visual Deprivation on the Neural Bases of Auditory Processing. Journal of Neuroscience, 36(5), 1620–1630. https://doi.org/10.1523/JNEUROSCI.2559-15.2016

Hötting, K., & Röder, B. (2004). Hearing cheats touch, but less in congenitally blind than in sighted individuals. Psychological Science, 15(1), 60–64

Huber, E., Chang, K., Alvarez, I., Hundle, A., Bridge, H., & Fine, I. (2019). Early blindness shapes cortical representations of auditory frequency within auditory cortex. The Journal of Neuroscience, 2896–18. https://doi.org/10.1523/JNEUROSCI.2896-18.2019

Jonas, C., Spiller, M. J., Hibbard, P. B., & Proulx, M. (2017). Introduction to the Special Issue on Individual Differences in Multisensory Perception: an Overview. Multisensory Research, 30(6), 461–466. https://doi.org/10.1163/22134808-00002594

Jones, E. G., & Powell, T. P. (1970). An anatomical study of converging sensory pathways within the cerebral cortex of the monkey. Brain, 93(4), 793–820. Retrieved from http://www.ncbi.nlm.nih.gov/pubmed/4992433

Kolarik, A. J., Cirstea, S., & Pardhan, S. (2013). Evidence for enhanced discrimination of virtual auditory distance among blind listeners using level and direct-to-reverberant cues. Exp. Brain Res, 224(4), 623–633. https://doi.org/10.1007/s00221-012-3340-0

Luo, Y. H. L., & da Cruz, L. (2016). The Argus® II Retinal Prosthesis System. Progress in Retinal and Eye Research, 50, 89–107. https://doi.org/10.1016/j.preteyeres.2015.09.003

Ma, W. J., Beck, J. M., Latham, P. E., & Pouget, A. (2006). Bayesian inference with probabilistic population codes. Nature Neuroscience, 9(11), 1432–1438. https://doi.org/10.1038/nn1790

Meijer, P. B. L. (1992). An experimental system for auditory image representations. In IEEE Transactions on Biomedical Engineering (Vol. 39, pp. 112–121). https://doi.org/10.1109/10.121642

Murray, M. M., Thelen, A., Ionta, S., & Wallace, M. T. (2018). Contributions of intraindividual and interindividual differences to multisensory processes. Journal of Cognitive Neuroscience, 31(3), 360–376. https://doi.org/10.1162/jocn_a_01246

Nagai, Y. (2019). Predictive learning: Its key role in early cognitive development. Philosophical Transactions of the Royal Society B: Biological Sciences, 374(1771), 20180030. https://doi.org/10.1098/rstb.2018.0030

Nardini, M., Bedford, R., & Mareschal, D. (2010). Fusion of visual cues is not mandatory in children. In Proceedings of the National Academy of Sciences (Vol. 107, pp. 17041–17046). https://doi.org/10.1073/pnas.1001699107

Nardini, M., Jones, P., Bedford, R., & Braddick, O. (2008). Development of cue integration in human navigation. Current Biology, 18(9), 689–693. https://doi.org/10.1016/j.cub.2008.04.021

Neil, P. A., Chee-Ruiter, C., Scheier, C., Lewkowicz, D. J., & Shimojo, S. (2006). Development of multisensory spatial integration and perception in humans. Developmental Science, 9(5), 454–464. https://doi.org/10.1111/j.1467-7687.2006.00512.x

Noppeney, U., Ostwald, D., & Werner, S. (2010). Perceptual decisions formed by accumulation of audiovisual evidence in prefrontal cortex. The Journal of Neuroscience : The Official Journal of the Society for Neuroscience, 30(21), 7434–7446. https://doi.org/10.1523/JNEUROSCI.0455-10.2010

Norman, J. F., & Bartholomew, A. N. (2011). Blindness enhances tactile acuity and haptic 3-D shape discrimination. *Attention*, Perception, and Psychophysics, 73(7), 2323–2331. https://doi.org/10.3758/s13414-011-0160-4

Oldfield, R. C. (1971). The assessment and analysis of handedness: The Edinburgh inventory. Neuropsychologia, 9(1), 97–113. https://doi.org/10.1016/0028-3932(71)90067-4

Ortiz-Terán, L., Ortiz, T., Perez, D. L., Aragón, J. I., Diez, I., Pascual-Leone, A., & Sepulcre, J. (2016). Brain plasticity in blind subjects centralizes beyond the modal cortices. Frontiers in Systems Neuroscience, 10, 61. https://doi.org/10.3389/fnsys.2016.00061

Pasqualotto, A., Furlan, M., Proulx, M. J., & Sereno, M. I. (2018). Visual loss alters multisensory face maps in humans. Brain Structure and Function, 223(8), 3731–3738. https://doi.org/10.1007/s00429-018-1713-2

Pasqualotto, A., & Proulx, M. J. (2012). The role of visual experience for the neural basis of spatial cognition. Neuroscience & Biobehavioral Reviews, 36, 1179–1187. https://doi.org/10.1016/j.neubiorev.2012.01.008

Peterzell, D. H. (2016). Discovering Sensory Processes Using Individual Differences : A Review and Factor Analytic Manifesto. Electronic Imaging, 2016(16), 1–11. https://doi.org/10.2352/ISSN.2470-1173.2016.16HVEI-112

Petrini, K., Caradonna, A., Foster, C., Burgess, N., & Nardini, M. (2016). How vision and self-motion combine or compete during path reproduction changes with age. Scientific Reports, 6, 29163. https://doi.org/10.1038/srep29163

Petrini, K., Remark, A., Smith, L., & Nardini, M. (2014). When vision is not an option: Children’s integration of auditory and haptic information is suboptimal. Developmental Science, 17(3), 376–387. https://doi.org/10.1111/desc.12127

Putzar, L., Hötting, K., & Röder, B. (2010). Early visual deprivation affects the development of face recognition and of audio-visual speech perception. Restorative Neurology and Neuroscience, 28(2), 251–257. https://doi.org/10.3233/RNN-2010-0526

Röder, B., Teder-Sälejärvi, W., Sterr, A., Rösler, F., Hillyard, S. A., & Neville, H. J. (1999). Improved auditory spatial tuning in blind humans. Nature, 400(6740), 162–166. https://doi.org/10.1038/22106

Rohde, M., van Dam, L. C. J., & Ernst, M. (2016). Statistically Optimal Multisensory Cue Integration: A Practical Tutorial. Multisensory Research, 29(4–5), 279–317. Retrieved from http://www.ncbi.nlm.nih.gov/pubmed/29384605

Rohe, T., Ehlis, A.-C., & Noppeney, U. (2019). The neural dynamics of hierarchical Bayesian causal inference in multisensory perception. Nature Communications, 10(1), 1907. https://doi.org/10.1038/s41467-019-09664-2

Sadato, N., Pascual-Leone, A., Grafman, J., Ibañez, V., Deiber, M. P., Dold, G., & Hallett, M. (1996). Activation of the primary visual cortex by Braille reading in blind subjects. Nature, 380(6574), 526–528. https://doi.org/10.1038/380526a0

Scheller, M., Petrini, K., & Proulx, M. J. (2018). Perception and Interactive Technology. In J. Wixted (Ed.), Stevens’ Handbook of Experimental Psychology and Cognitive Neuroscience (4th ed., pp. 1–50). John Wiley & Sons, Inc. https://doi.org/10.1002/9781119170174.epcn215

Sowell, E. R., Peterson, B. S., Thompson, P. M., Welcome, S. E., Henkenius, A. L., & Toga, A. W. (2003). Mapping cortical change across the human life span. Nature Neuroscience, 6(3), 309–315. https://doi.org/10.1038/nn1008

Stanley, B. M., Chen, Y.-C., Lewis, T. L., Maurer, D., & Shore, D. I. (2019). Developmental changes in the perception of audiotactile simultaneity. Journal of Experimental Child Psychology, 183, 208–221. https://doi.org/10.1016/j.jecp.2019.02.006

Vercillo, T., Burr, D., & Gori, M. (2016). Early visual deprivation severely compromises the auditory sense of space in congenitally blind children. Developmental Psychology, 52(6), 847–853. https://doi.org/10.1037/dev0000103

Vercillo, T., Milne, J. L., Gori, M., & Goodale, M. A. (2015). Enhanced auditory spatial localization in blind echolocators. Neuropsychologia, 67, 35–40. https://doi.org/10.1016/j.neuropsychologia.2014.12.001

Voss, P., Gougoux, F., Lassonde, M., Zatorre, R. J., & Lepore, F. (2006). A positron emission tomography study during auditory localization by late-onset blind individuals. NeuroReport, 17(4), 383–388. https://doi.org/10.1097/01.wnr.0000204983.21748.2d

Voss, P., Lassonde, M., Gougoux, F., Fortin, M., Guillemot, J. P., & Lepore, F. (2004). Early- and late-onset blind individuals show supra-normal auditory abilities in far-space. Current Biology, 14(19), 1734–1738. https://doi.org/10.1016/j.cub.2004.09.051

Wan, C. Y., Wood, A. G., Reutens, D. C., & Wilson, S. J. (2010). Early but not late-blindness leads to enhanced auditory perception. Neuropsychologia, 48(1), 344–348. https://doi.org/10.1016/j.neuropsychologia.2009.08.016

World Health Organization. (2018). International statistical classification of diseases and related health problems (11th Revision). 9D90 Vision impairment including blindness. Retrieved from https://icd.who.int/browse11/l-m/en

Zwiers, M. P., Van Opstal, A. J., & Cruysberg, J. R. (2001). A spatial hearing deficit in early-blind humans. The Journal of Neuroscience : The Official Journal of the Society for Neuroscience, 21(9), RC142: 1-5. Retrieved from http://www.ncbi.nlm.nih.gov/pubmed/11312316

## References

Brainard, D. H. (1997). The Psychophysics Toolbox. Spatial Vision, 10(4), 433–436. https://doi.org/10.1163/156856897X00357

Fischer, J., & Whitney, D. (2014). Serial dependence in visual perception. Nature Neuroscience, 17(5), 738–743. https://doi.org/10.1038/nn.3689

Grassi, M. (2005). Do we hear size or sound? Balls dropped on plates. Perception and Psychophysics, 67(2), 274–284. https://doi.org/10.3758/BF03206491

Kavšek, M., & Granrud, C. E. (2012). Children’s and adults’ size estimates at near and far distances: A test of the perceptual learning theory of size constancy development. I-Perception, 3(7), 459–466. https://doi.org/10.1068/i0530

Nardini, M., Begus, K., & Mareschal, D. (2013). Multisensory uncertainty reduction for hand localization in children and adults. Journal of Experimental Psychology: Human Perception and Performance, 39(3), 773–787. https://doi.org/10.1037/a0030719

Pelli, D. G. (1997). The VideoToolbox software for visual psychophysics: transforming numbers into movies. Spatial Vision, 10(4), 437–442. https://doi.org/10.1163/156856897X00366

Prins, N. (2013). The psi-marginal adaptive method: How to give nuisance parameters the attention they deserve (no more, no less). Journal of Vision, 13(7), 3. https://doi.org/10.1167/13.7.3

Prins, N., & Kingdom, F. A. A. (2018). Applying the Model-Comparison Approach to Test Specific Research Hypotheses in Psychophysical Research Using the Palamedes Toolbox. Frontiers in Psychology, 9, 1250. https://doi.org/10.3389/fpsyg.2018.01250

